# Comprehensive double-mutant analysis of the *Bacillus subtilis* envelope using double-CRISPRi

**DOI:** 10.1101/2024.08.14.608006

**Authors:** Byoung-Mo Koo, Horia Todor, Jiawei Sun, Jordi van Gestel, John S. Hawkins, Cameron C. Hearne, Amy B. Banta, Kerwyn Casey Huang, Jason M. Peters, Carol A. Gross

**Affiliations:** Department of Microbiology and Immunology, University of California, San Francisco, San Francisco, CA, USA; Department of Bioengineering, Stanford University, Stanford, CA, USA; Pharmaceutical Sciences Division, School of Pharmacy, University of Wisconsin-Madison, Madison, Wisconsin, USA; Department of Microbiology and Immunology, Stanford University School of Medicine, Stanford, CA, USA; Chan Zuckerberg Biohub, San Francisco, CA, USA; Department of Cell and Tissue Biology, University of California, San Francisco, San Francisco, California, USA; California Institute of Quantitative Biology, University of California, San Francisco, San Francisco, CA, USA

**Author notes:** Contributed equally (Byoung-Mo Koo, Horia Todor).

**Keywords:** Genetic interaction, Double-CRISPRi, Cell envelope, *mbl*, *mreB*, Cell division

## Abstract

Understanding bacterial gene function remains a major biological challenge. Double-mutant genetic interaction (GI) analysis addresses this challenge by uncovering the functional partners of targeted genes, allowing us to associate genes of unknown function with novel pathways and unravel connections between well-studied pathways, but is difficult to implement at the genome-scale. Here, we develop and use double-CRISPRi to systematically quantify genetic interactions at scale in the *Bacillus subtilis* envelope, including essential genes. We discover > 1000 known and novel genetic interactions. Our analysis pipeline and experimental follow-ups reveal the distinct roles of paralogous genes such as the *mreB* and *mbl* actin homologs, and identify new genes involved in the well-studied process of cell division. Overall, our study provides valuable insights into gene function and demonstrates the utility of double-CRISPRi for high-throughput dissection of bacterial gene networks, providing a blueprint for future studies in diverse bacterial species.

## INTRODUCTION

The field of genetics has been built on deducing gene functions by associating gene disruptions with phenotypes. The ability to investigate the phenotypes of gene disruption mutants in bacteria at genome-scale using high throughput techniques such as transposon insertion libraries (Price et al., 2018; van Opijnen et al., 2009), single-gene deletion collections (Koo et al., 2017; Nichols et al., 2011), and CRISPR interference (CRISPRi) libraries (Liu et al., 2021; Peters et al., 2016; Wang et al., 2018; Yao et al., 2020) has dramatically advanced the pace of discovery of gene functions and enabled unbiased discovery of functional partners through shared phenotypes. Such chemical-genomic studies have predicted functions for thousands of previously uncharacterized or poorly characterized genes and revealed new connections between cellular pathways. Nonetheless, studies with hundreds of distinct conditions have failed to identify any phenotypes for a significant fraction of genes (∼30%) even in the best studied model organisms such as *Escherichia coli* (Nichols et al., 2011). Determining the functions of these genes, many of which are broadly conserved, is an outstanding problem.

Genetic interaction (GI) mapping, a cornerstone of classical genetic approaches (Mani et al., 2008), compares the phenotypes of double-deletion mutant strains to the sum of their single knockout phenotypes. Differences from the null expectation are indicative of genetic interactions (GIs), which can reveal the interacting partners of a gene and uncover phenotypes (e.g. essentiality) for partially redundant gene pairs. The power of this approach has been demonstrated by numerous studies that mapped the GIs between a single gene and the rest of the genome (1 × all) to discover novel protein functions like undecaprenyl flippases (Sit et al., 2023), peptidoglycan (PG) polymerase regulators (Paradis-Bleau et al., 2010), and PG hydrolase co-factors (Brunet et al., 2019). Despite the utility of double-mutant analyses, genome-scale GI screens have thus far been executed only in the yeast *Saccharomyces cerevisiae* (Costanzo et al., 2016), for which automated construction and analysis of >20 million double mutants revealed overall construction principles of the cell. The bottleneck to general use of large-scale GI analysis is that screening requires constructing double mutants in large pools and then determining the identity of both affected genes, even when they are distant from each other on the chromosome. These challenges can be overcome using CRISPRi. Two genes can be transcriptionally repressed by adjacently encoded sgRNAs and the sgRNAs can be identified and enumerated via sequencing. A CRISPRi-based GI approach has been demonstrated by interrogating 222,784 double knockdown strains (472 × 472 genes) in mammalian cells (Horlbeck et al., 2018) but has not been applied to bacteria at genome-scale.

Here, we develop double-CRISPRi technology in the model Gram-positive bacterium *Bacillus subtilis* and use it to perform a genome-scale GI screen of envelope genes. We chose to focus on the Gram-positive cell envelope, which is composed of the inner membrane (IM), the peptidoglycan (PG) cell wall, and associated molecules such as teichoic acids (TA) (Silhavy et al., 2010), for several reasons. First, the envelope is responsible for cellular integrity, elongation, and division, and for mediating environmental, pathogenic, and symbiotic interactions. Second, since the envelope is the target of many antibiotics (Jordan et al., 2008; Page and Walker, 2021; Sarkar et al., 2017), identification of synthetic-lethal gene pairs can aid the design of synergistic antibiotic therapies. Third, envelope processes are difficult to reconstitute biochemically, as they have numerous components and often function across many length scales (Rohde, 2019; Typas et al., 2011), making genetic dissection paramount. However, the partial redundancy of envelope-function genes necessary to ensure robust growth across conditions has complicated this genetic dissection (McPherson and Popham, 2003; Straume et al., 2021; Thomaides et al., 2007). Finally, despite intense study over decades, the cell envelope still contains the highest fraction of proteins of unknown function (Hu et al., 2009; Pedreira et al., 2022).

Our experiments identified >1000 known and novel positive and negative GIs. By combining our screen with follow-up experiments including live cell microscopy, we uncover links between diverse envelope processes that expand our understanding of the Gram-positive cell envelope and provide a valuable resource and discovery tool for the research community. Our study provides a natural stepping-stone to genome-wide screens, which remain technically and financially challenging due to their size (∼4,000 x ∼4,000 genes = ∼16 million total strains).

## RESULTS AND DISCUSSION

### Construction of a chromosomally encoded double-CRISPRi library

Our double-CRISPRi system is based on a xylose-inducible, chromosomally-integrated single-gene knockdown CRISPRi system (Hawkins et al., 2020; Peters et al., 2016). In double-CRISPRi, two sgRNAs targeting different genes are placed adjacent to each other on the chromosome such that double knockdown strain abundances can be quantified from paired-end sequencing reads (Figure 1A). Because loss of repeated DNA sequences is generally high (∼10^-4^/generation in *B. subtilis*, and potentially higher in certain growth conditions or mutants), we took steps to minimize sgRNA loss via loop-out and recombination either during the experiment or during DNA sequencing (Reis et al., 2019). We used different but equally strong constitutive promoters, terminators, and sgRNA scaffolds for the adjacently encoded sgRNAs. In addition to these changes at the sgRNA locus, we also replaced the erythromycin-resistance marker adjacent to *dcas9* with a kanamycin-resistance marker flanked by *lox* sequences (Figures 1A, S1A and S1B). This change eliminated ribosome methylation by the erythromycin-resistance protein, which can affect bacterial physiology (Gupta et al., 2013), and enables the removal and reuse of the kanamycin marker for strain construction in follow-up studies (Koo et al., 2017; Yan et al., 2008).

**Figure 1.**
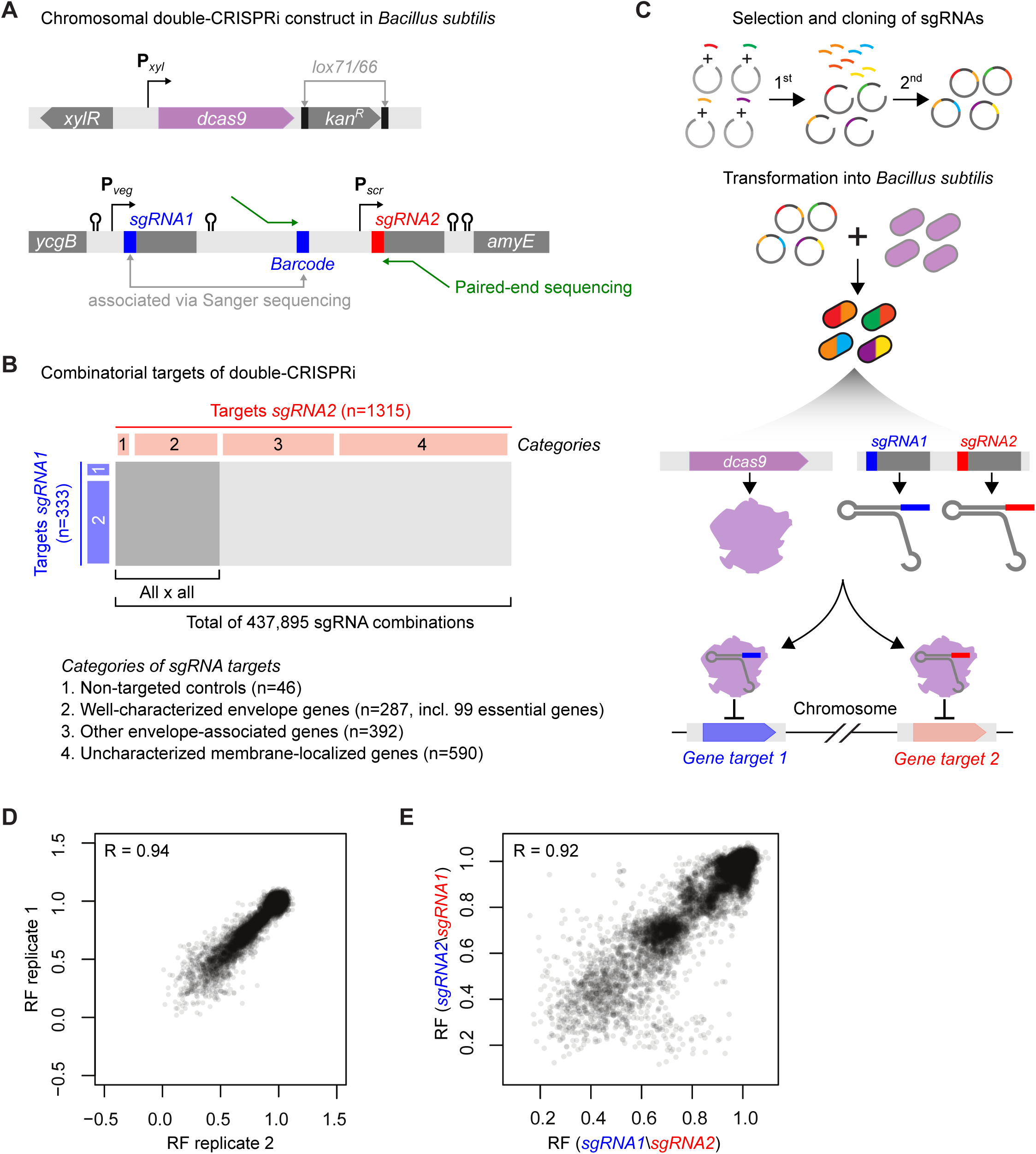
A large-scale inducible, chromsomally-integrated double-CRISPRi system in *B. subtilis*. **A)** The structure of the *dcas9* and sgRNA loci. Upper: inducible *dcas9* expression system. The Kanamycin-resistant gene is flanked by *lox71/66* sites which can be removed by Cre recombinase. Lower: Double gene knockdown system used in this study. Two constitutive promoters of similar strength, P*_veg_* and P*_scr_*, transcribe the first and second sgRNAs, respectively. A 26bp barcode is inserted between the first and second sgRNAs and associated with the first sgRNA via Sanger sequencing to facilitate sequencing-based identification. Four different but equivalent transcriptional terminators ensure independent transcription of each sgRNA. The final *B. subtilis* strain contains a xylose-inducible *dcas9* gene at the *lacA* locus and the two sgRNAs at the *amyE* locus. **B)** Identity of the envelope gene pairs assayed by double-CRISPRi (Table S1). **C)** Schematic of the Double-CRISPRi library construction method. The DNA fragments containing the sgRNAs and their associated random barcodes were cloned individually at the first position. These plasmids were then pooled and the sgRNAs at the second position were cloned using a pooled approach (Methods). **D)** Correlation between RF of two representative biological replicates. **E)** Correlation between RFs of the same gene pairs with sgRNAs in the opposite order (sgRNA1\sgRNA2 vs sgRNA2\sgRNA1).

Our screen targeted two sets of envelope function genes (Figure 1B). The first set consists of well-characterized envelope genes, select essential genes, and non-targeting controls (333 total; categories 1,2; Figure 1B). Essential genes were targeted by mismatched sgRNAs (Hawkins et al., 2020) that produce mild knockdown and moderate growth defects, to enable the quantification of both positive and negative GIs. The second set consists of poorly characterized membrane-localized or envelope-associated genes (982 total; categories 3,4; Figure 1B and Table S1). By using these two sets, we can identify new connections between well-studied pathways and associate poorly characterized or peripheral envelope genes with established pathways.

To construct the library, we first cloned the sgRNA targeting each gene in the first set and a random barcode into the first sgRNA position, and then associated the two via Sanger sequencing (Figure 1C). Most of the sgRNAs (316/333, 95%) were successfully cloned. We next cloned sgRNAs from both sets (1315 total) as a pool into the second sgRNA position, resulting in a library querying 415,540 potential gene-gene interactions (316×1315). 93% of the potential double-CRISPRi strains were successfully constructed, and our cloning process resulted in a tight distribution of strain abundances, with ∼90% of strains within 10-fold of the median (Figure S1C). This high-quality library facilitates high-throughput screening of envelope gene GIs and provides a blueprint for double-CRISPRi library construction targeting diverse gene sets.

### Double-CRISPRi identifies high-quality GIs

dCas9 was induced in cells undergoing exponential growth (maintained via back dilution). Cells were sampled immediately before dCas9 induction and after 10 doublings post induction, as well as at several other time points (Figure S2). The relative fitness (RF; (Hawkins et al., 2020; Kampmann et al., 2013; Rest et al., 2013)) of each strain was calculated by comparing its relative abundance at the start and end of each experiment (Methods; Table S2). Libraries were sequenced to high read depth (median read depth per strain ∼ 500) to enable accurate RF measurements of slow-growing strains. The RF of individual double-CRISPRi strains was highly correlated across replicates (Pearson’s *r* ∼ 0.94, Figures 1D and S3A) and with previously published single-CRISPRi experiments (Figure S3B) (Hawkins et al., 2020). Importantly, RFs were highly correlated between strains containing the same two sgRNAs in the opposite order (sgRNA1\sgRNA2 versus sgRNA2\sgRNA1; Pearson’s *r* ∼ 0.92; Figure 1E), despite differences in the promoters, sgRNA scaffolds, and terminators driving expression of the two sgRNAs.

To quantify GIs, we compared the RF of each double-knockdown strain to the RF of its two parent strains using an approach that conveys information about both the strength and statistical significance of a GI (modified from (Collins et al., 2006) Methods; Figure S4). Positive GI scores occur when a double knockdown strain grows better than expected based on the growth defects of its parent strains (e.g., one gene is a suppressor of the other). Negative GI scores occur when a double knockdown strain grows worse than expected based on the growth defects of its parent strains (e.g., the genes are synthetic sick/lethal). The final data set (Table S3) was comprised of GI scores for ∼291,000 double-mutant strains passing a set of stringent quality control standards (e.g., minimum read depth, multiple replicates, no correlation to sgRNA sequence; see Methods). Consistent with the idea that GIs are rare (Hartman et al., 2001), most GI scores were ∼0 (Figure 2A). The GI scores of gene pairs with a strong (|GI score|>3) or significant (|GI score|>2) GI in at least one replicate were highly correlated between replicates (*r ∼* 0.79 for 2400 interactions with |GI score|>3; *r* ∼ 0.59 for 15,000 interactions with |GI score|>2; Figure S5A). GI scores were also correlated between genes within the 22 operons with very strong (|GI score|>5) GIs (median within operon Pearson’s *r* ∼ 0.25, Figure S5B).

**Figure 2.**
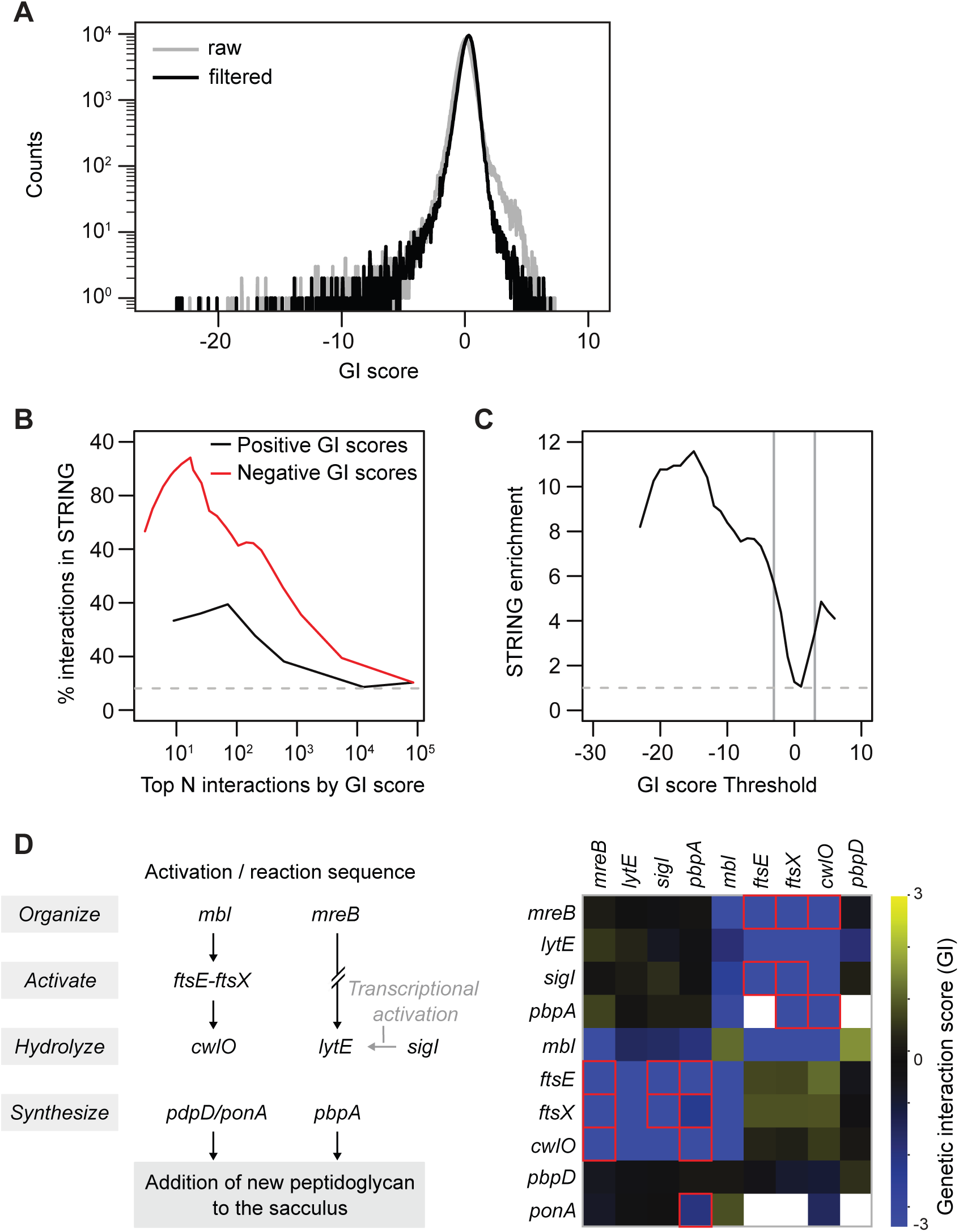
Double-CRISPRi accurately and sensitively identifies genetic interactions. **A)** The distribution of GI scores in the library after 10 cell doublings with (black) and without (gray) filtering (Methods). Most GI scores are near zero, consistent with the hypothesis that most gene pairs do not interact. **B)** Gene pairs with strongly positive (black) or negative (red) GI scores have a high proportion of gene pairs with evidence of physical or genetic interactions from the STRING database. **C)** Gene pairs with strongly positive or negative GI scores are enriched in gene pairs with evidence of physical or genetic interactions from the STRING database. Gray lines indicate the GI score threshold at +/-3. **D)** Double-CRISPRi recapitulates known interactions between two cell wall hydrolysis activation pathways. Left) Schematic of the two cell wall hydrolysis/synthesis pathways. SigI is a transcriptional activator of *lytE*, which is indicated by a gray arrow. Right) Heatmap of GI between genes involved in these pathways. Red boxes denote novel interactions identified in this screen.

The large knowledge base of interactions from previous envelope-focused studies enabled us to gauge whether our quantification of GIs accurately identified known interactions. First, we found that gene pairs with high absolute GI scores (both positive and negative) were enriched in all interactions documented in the STRING database (Szklarczyk et al., 2023) (GI score>3: ∼3.4-fold enriched, 56/202 interactions in STRING; GI score<-3: ∼5.6-fold enriched, 268/587 interactions in STRING; Figure 2B & 2C). The STRING database contains known and predicted protein-protein interactions (PPIs) derived from physical, functional, and genomic associations. Second, our data set recapitulated well-characterized synthetic lethal phenotypes, identified novel GIs that are consistent with and extend known biology, and identified novel interactions in these pathways. For example, we identified the known synthetic lethality between the two cell-wall hydrolases, *cwlO* and *lytE* (Bisicchia et al., 2007), and also identified the novel but expected negative GIs between their activation pathways (Dominguez-Cuevas et al., 2013), as well as unexpected negative GIs between hydrolases and PG synthases (e.g. *pbpA*/cwlO and *pbpA*/*ftsEX*) that suggest an intimate connection between PG hydrolysis and synthesis (Figure 2D). Importantly, while known and expected GIs are significantly enriched in our data set, we also found many high-confidence novel interactions that further illuminate cell envelope function. These novel interactions include well-studied genes such as the essential actin homologs *mreB* and *mbl* (Figure S5C), as well as less studied genes, indicating that our data set can function as an engine for discovery (Table S3).

### Correlated profiles of GIs identify interacting genes

A gene’s pattern of GIs can be used to provide additional insight into its function by providing quantitative phenotypes that can be compared collectively to identify functionally related genes (Collins et al., 2007; Horlbeck et al., 2018), analogous to the implications of correlated chemical sensitivities in chemical genomics screens (Figure 3A) (Nichols et al., 2011; Peters et al., 2016; Shiver et al., 2016). Consistent with this idea and with analyses in yeast (Collins et al., 2007) and human cells (Horlbeck et al., 2018), we found that gene pairs with highly correlated GI profiles were enriched in previously discovered interactions (Pearson’s *r* > 0.5, ∼7.3-fold enriched, 141/300 interactions in STRING; Figure 3B). Hierarchical clustering of the matrix of GI score correlations distinguished cell division, cell-wall hydrolysis, and other envelope processes (Figure 3C).

**Figure 3.**
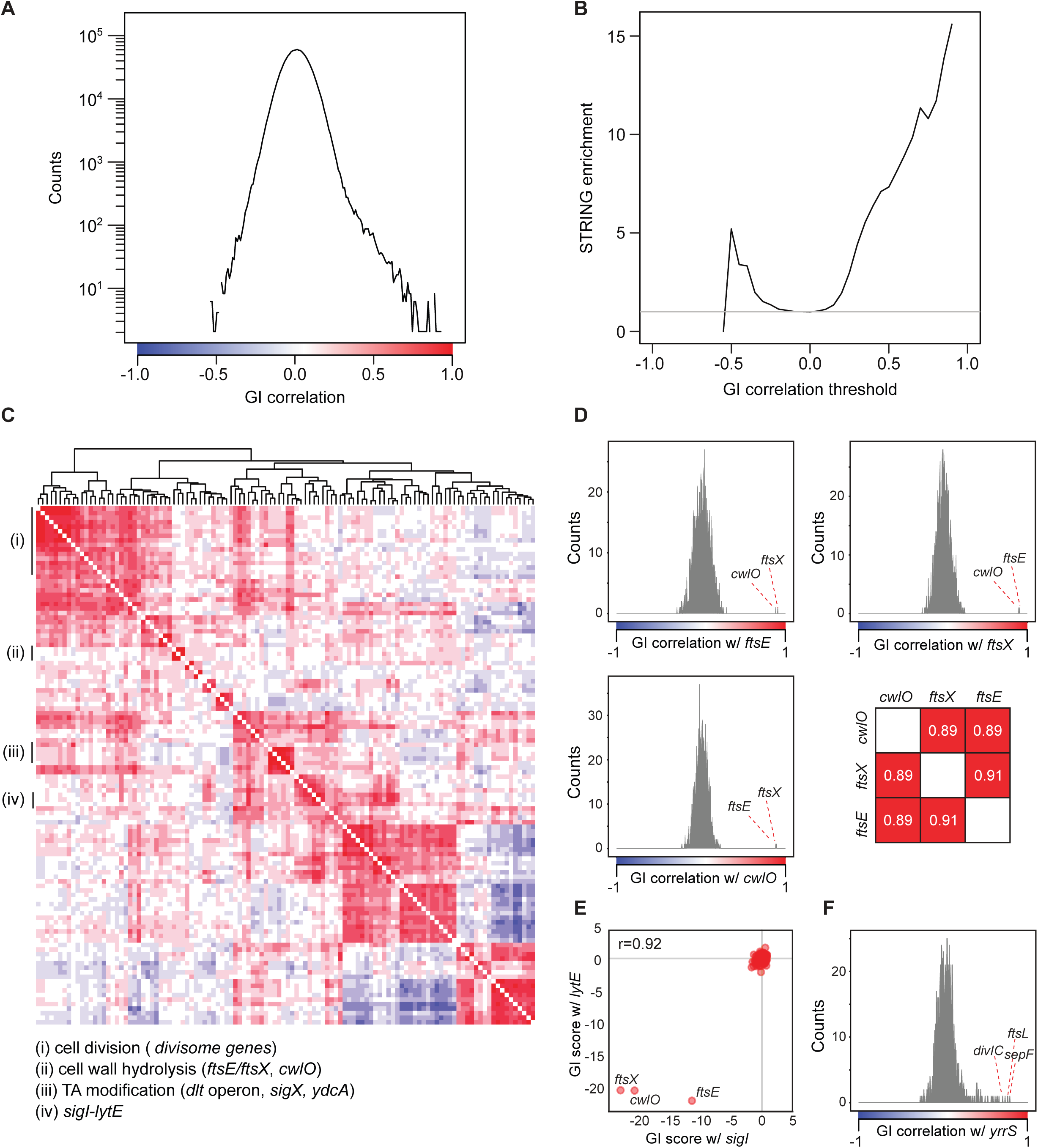
Correlated GI scores identify co-functioning gene pairs. **A)** The distribution of GI score correlations between sgRNAs at the second position. **B)** Correlated (and anti-correlated) gene pairs are enriched in those with evidence of physical or genetic interactions from the STRING database. **C)** Clustered heatmap of genes with at least one strong correlation (>0.5) reveals co-functioning genes. Colors indicate the Pearson correlation (Figure 3A). The uncharacterized gene *ydcA* was clustered with genes involved in TA modification, likely due to a polar effect (Supplementary Note 2). **D)** Histograms of the correlation scores of *ftsE*, *ftsX* and *cwlO* showing specific interactions between these three genes. **E)** GI scores for all strains with *lytE* (x-axis) and *sigI* (y-axis). **F)** A histogram of correlations between all genes and *yrrS* suggests a role for *yrrS* in cell division.

Further analysis revealed three biologically relevant reasons for highly correlated gene pairs. First, genes encoding proteins in the same pathway exhibited highly correlated GI profiles. For example, FtsE and FtsX are required for the activity of the CwlO peptidoglycan hydrolase (Meisner et al., 2013). *ftsE*, *ftsX*, and *cwlO* exhibited highly correlated GI profiles with each other (*r* > 0.88), but not with other genes (the next strongest correlation was <0.31, Figure 3D). Moreover, the three genes had no strong GIs with each other (highest |GI score|<1.3). Second, some sigma factors exhibited GI profiles highly correlated to that of genes in their regulon. For example, SigI has a small regulon that notably includes *lytE* (Ramaniuk et al., 2018); *sigI* and *lytE* profiles were highly correlated (*r*>0.92, Figure 3E). Finally, GI profiles were highly correlated among members of functional protein complexes, such as the divisome (Halbedel and Lewis, 2019) (Figure 3C). These correlations suggest a novel role for the poorly characterized gene *yrrS* in cell division, based on strong correlations to the GI profile of known cell division genes such as *sepF*, *ftsL*, *and divIC* (*r*>0.7, Figure 3F). Taken together, these data indicate that correlated GI score profiles provide additional insight into the function of envelope genes.

### GI analysis reveals distinct functions of paralogous genes

Duplication and divergence of genes are major drivers of evolution, and as a result, paralogous genes are common in bacteria (Hernandez-Plaza et al., 2023). However, our understanding of the shared and distinct functions of these genes is often incomplete. Since the GI profiles of paralogous genes can illuminate the degree of functional divergence, we examined the GIs of three pairs of partially redundant paralogous genes: the undecaprenyl pyrophosphate phosphatases *bcrC* and *uppP*, the lipoteichoic acid (LTA) synthases *ltaS* and *yfnI*, and the actin homologs *mreB* and *mbl.* In each case, knockdown of both paralogs led to a strong GI, confirming their redundant functionality (Figure 4) (Jones et al., 2001; Radeck et al., 2017; Wormann et al., 2011). However, each paralog exhibited some distinct GIs in our screen, providing clues to their specialized function.

**Figure 4.**
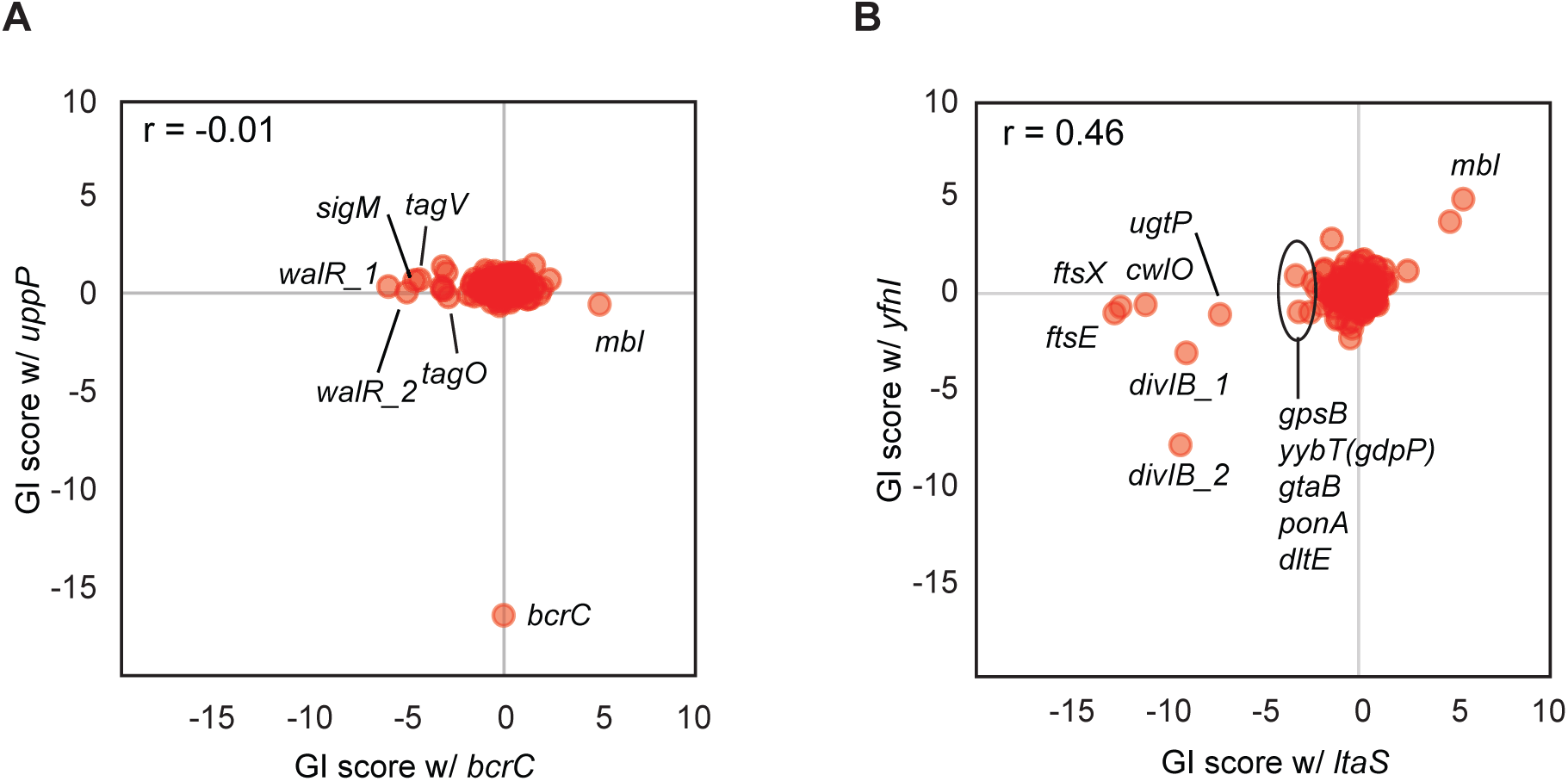
Distinct functions of paralogous genes. **A)** GI scores for strains containing *bcrC* or *uppP* targeting sgRNAs. *walR_1* and *walR_2* represent two different sgRNAs targeting *walR*. **B)** GI scores for the strains containing *ltaS* or *yfnI* targeting sgRNAs. *divIB_1* and *divIB_2* represent two different sgRNAs targeting *divIB*.

*bcrC* and *uppP* each encode a key enzyme in the lipid II cycle (undecaprenyl pyrophosphate phosphatase) and have approximately equivalent transcript levels (Radeck et al., 2017). BcrC is proposed to be the major enzyme in *B. subtilis* (Inaoka and Ochi, 2012), and consistent with this designation, a Δ*bcrC* but not a Δ*uppP* mutant exhibited a slow-growth phenotype (Radeck et al., 2016). As expected based on these observations, our data indicated that *uppP* has only one strong GI (synthetic lethal with *bcrC* (Zhao et al., 2016)), whereas *bcrC* has many strong GIs, including a strong negative interaction with *sigM* (Figure 4A). Since *sigM* becomes essential under undecaprenyl phosphate (Und-P)-limiting conditions (Roney and Rudner, 2024), the strong negative GI between *bcrC* and *sigM* suggests that BcrC depletion significantly reduces Und-P levels. Other GIs with *bcrC* motivate additional study. For example, *bcrC* has strong negative interactions with the most upstream gene involved in wall teichoic acid (WTA) synthesis, *tagO,* and the phosphotransferase gene for WTA attachment, *tagV* (Figure 4A). These negative interactions could result from disruption of the lipid II cycle, as these two enzymes use or produce Und-P after their catalytic reactions (Gale et al., 2017; Soldo et al., 2002). Interestingly, in *E. coli*, the roles of the two Und-Pases are reversed: *uppP* is responsible for 75% of undecaprenyl pyrophosphate phosphatase activity while *bcrC* is considered a minor enzyme (El Ghachi et al., 2004).

### *ltaS* and *yfnI* are partially redundant paralogs involved in LTA synthesis

Compared to *ltaS*, *yfnI* (which is activated by stress) encodes an enzyme that produces longer LTAs (Jervis et al., 2007; Wormann et al., 2011). Both *ltaS* and *yfnI* exhibited strong negative GIs with *divIB*, an essential member of divisome (Figure 4B). However, *ltaS* but not *yfnI* exhibited strong negative interactions with *ftsEX* and *cwlO,* suggesting that the different LTA polymers produced by these paralogs have differential effects on the PG elongation machinery (Figure 4B). Moreover, knockdown of *ltaS* but not *yfnI* showed weak but consistent negative interactions with the *dlt* genes (Figure 4B), a phenotype we validated using double deletion mutants (Figure S6). In the absence of *ltaS*, all LTAs are of the *yfnI* type. The *dlt* genes are involved in D-alanylation of TAs (Perego et al., 1995), suggesting that LTAs synthesized by YfnI but not those synthesized by LtaS require D-alanylation for full functionality.

MreB and Mbl are well-studied essential paralogs that function in cell shape determination through regulation of cell-wall elongation (Jones et al., 2001). Whereas *mreB* is almost universally conserved in rod-shaped bacteria, additional *mreB* homologs such as *mbl* are found exclusively in Gram-positive phyla (Takahashi et al., 2020). *mbl* or *mreB* can be deleted in the presence of excess Mg^2+^, which stabilizes the cell envelope and inhibits the activity of LytE and perhaps other PG hydrolases, and both knockout strains exhibit morphological defects (Formstone and Errington, 2005; Schirner and Errington, 2009; Tesson et al., 2022). As expected, *mreB* and *mbl* exhibited a strong negative GI in our screen (Figures 2D and S5C). *mreB* and *mbl* both had negative GIs with *ftsE*, *ftsX*, and *cwlO*, confirming their synergistic role in guiding the elongation machinery and controlling the activity of cell-wall hydrolases (Figures 2D and S5C). Strikingly, although we identified many positive (suppressive) GIs for *mbl*, including known suppressors such as *ltaS* (Schirner et al., 2009), we did not identify any suppressors for *mreB* (Figures 5A, blue quadrant; and S5C), an observation we follow up in subsequent sections. Taken together, these data suggest that GI analysis can disentangle the shared and unique functions of partially redundant genes.

**Figure 5.**
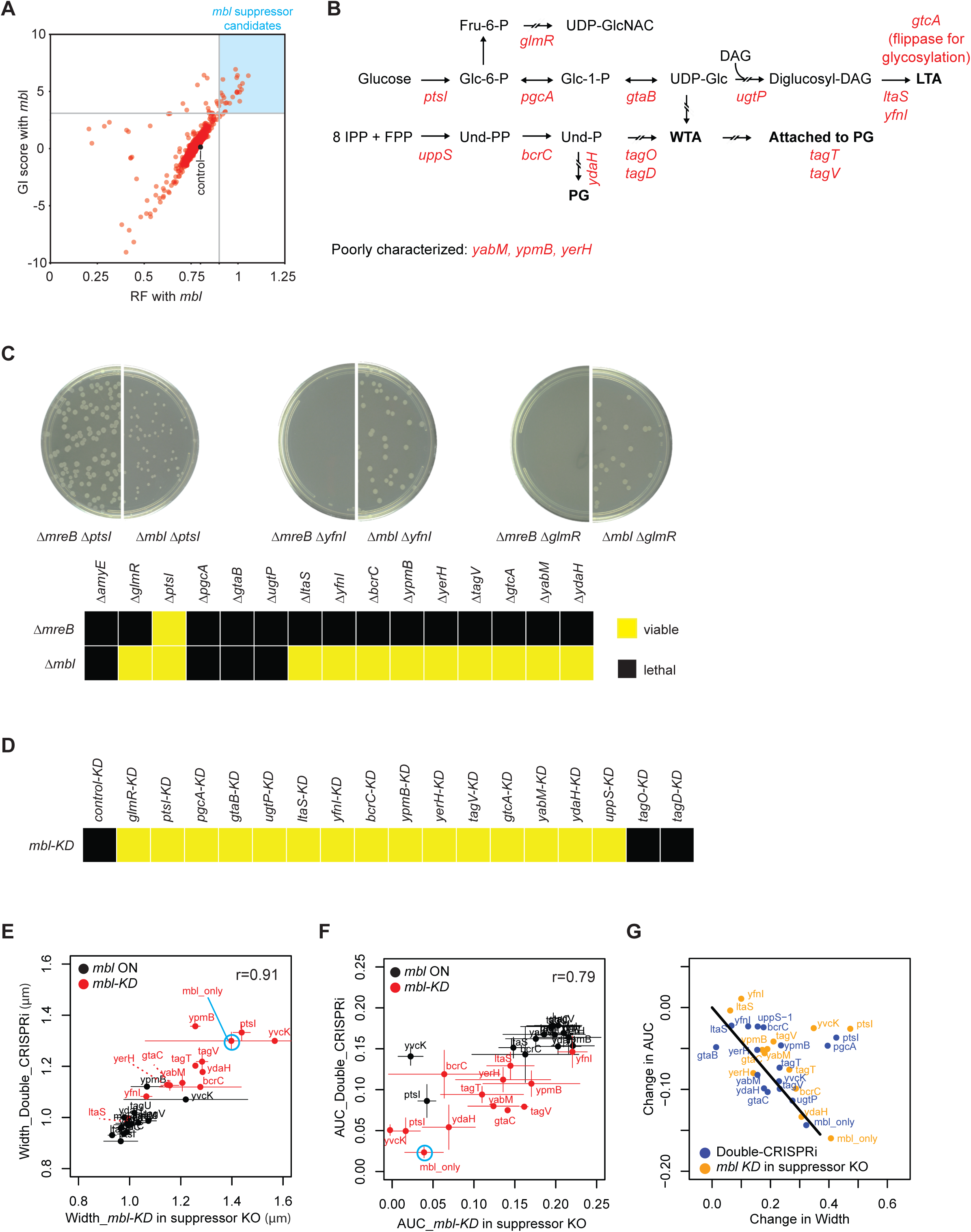
Genetic interactions disentangle the roles of *mbl* and *mreB* in cell envelope homeostasis. **A)** *mbl* suppressors for follow-up were selected by looking for *mbl* double-CRISPRi strains with RF > 0.9 and a GI score > 3. The RF of *mbl*-control strains was ∼0.78 (gray dot). **B)** *mbl* suppressors were involved in various aspects of WTA, LTA, and PG synthesis. *glmR* is a positive regulator for UDP-GlcNAC synthesis. UDP-GlcNAC is a precursor for PG and other metabolites. **C)** Construction of *mbl* and *mreB* deletions in *mbl* specific and common suppressor background without high concentration of Mg^2+^. *mbl::kan* and *mreB::kan* fragments were transformed into each genetic background strain and incubated for 16 hours. Upper: Plate images of the transformation of *mbl::kan* and *mreB::kan* into Δ*ptsI*, Δ*yfnI*, and Δ*glmR* strains. Lower: Viability of the double mutants constructed by the transformation of *mbl::kan* and *mreB::kan* into *mbl* suppressor deletion strains. **D)** Viability of the individual double-CRISPRi knockdown strains of all mbl/suppressor pairs. Viable knockdown pairs are indicated by yellow squares as shown in Figure 5C. **E)** Growth rescue and cell width change after *mbl* and suppressor double CRISPRi knockdown. The area under the curve (AUC) indicates growth. The open circle indicates cells before *mbl* knockdown and the closed circle indicates cells after *mbl* knockdown. **F)** Growth rescue and cell width change after *mbl* knockdown in suppressor deletion background. **G)** Correlation between suppressor gene deletion and suppressor gene knockdown for cell width changes upon *mbl* knockdown. **Abbreviations**: Fru-6-P: fructose-6-phosphate; UDP-GlcNAC: UDP-N-acetylglucosamine; Glc-6-P: glucose-6-phosphate; Glc-1-P:glucose-1-phosphate; UDP-Glc: UDP-glucose; DAG: diacylglycerol; IPP: isopentenyl pyrophosphate; FPP: farnesyl pyrophosphate; Und-PP: undecaprenyl pyrophosphate: Und-P: undecaprenyl phosphate; PG: peptidoglycan.

### Dissecting the role of *mbl* in TA synthesis

Our double-CRISPRi analysis revealed a striking difference in the GI profiles of *mbl* (many strong positive GIs) and *mreB* (no strong positive GIs). Genes in several processes had positive (suppressive) GIs with *mbl*. First, disrupting genes involved in LTA synthesis rescued *mbl* knockdown (Figure 5B, upper). This group includes the major and minor LTA synthases (*ltaS*, *yfnI*), the LTA glycosylation protein (*gtcA*), and genes involved in TA precursor synthesis (*pgcA*, *gtaB*, *ugtP*). Second, many genes involved in WTA synthesis (*tagO* and *tagD*), lipid carrier cycling (*uppS*, *bcrC*, and *ydaH*), and attachment (*tagT* and *tagV*) had positive GIs with *mbl*, suggesting a previously unrecognized role for WTAs in regulating cell elongation (Figure 5B lower). Finally, genes involved in sugar metabolism (*ptsI* and *glmR*), and several poorly characterized genes (*ypmB*, *yerH*, *and yabM*) had positive GIs with *mbl*. Strikingly, none of these interactions were shared with *mreB*. Two technical considerations could potentially explain this observation. First, *mreB* was targeted by a mismatched sgRNA (partial knockdown, mild phenotype) while *mbl* was targeted by a fully complementary sgRNA (full knockdown, strong phenotype). Second, *mreB* is located in an operon that includes *mreC*, *mreD*, and other cell division genes, which may influence its GIs.

To validate the GI profiles of *mreB* and *mbl* using an orthogonal approach, we tested whether *mbl* or *mreB* deletion alleles could be transformed into strains carrying a deletion of each putative *mbl* suppressor gene identified in our double-CRISPRi screen (Figure 5C). In our growth/media conditions, we found that an *mbl* deletion allele could be transformed into almost all suppressor gene deletion strains. In stark contrast, the *mreB* deletion allele could not be transformed into any of the strains tested except for Δ*ptsI*, a known *mreB* suppressor (Kawai et al., 2009) (Figure 5C). Additional evidence for the differential roles of *mbl* and *mreB* is their interaction with *glmR*: whereas a Δ*glmR* suppresses *mbl* essentiality (Figure 5C), *glmR* must be overexpressed to suppresses *mreB* essentiality (Foulquier et al., 2011). Taken together, these data validate the results of our double-CRISPRi screen and greatly expand the universe of *mbl* interacting processes, adding both WTA synthesis and genes of unknown function. Notably, although double deletion strains of *mbl* and suppressor genes were viable in exponential phase, survival into stationary phase required activation of PG synthesis systems (Supplementary Note 1).

Knockout or CRISPRi knockdown of *mbl* results in cell widening prior to lysis (Peters et al., 2016; Schirner et al., 2009). Interestingly, knockout of *ltaS* restores both wild-type growth and morphology to *mbl-*disrupted cells (Schirner et al., 2009). We therefore tested whether both phenotypes were rescued by the additional suppressors we identified, including those that could not be reconstructed as double knockouts. We reconstructed all *mbl*/suppressor pairs as individual double-CRISPRi knockdown strains, including the essential gene suppressors (*tagO*, *tagD*, *uppS*) and LTA precursor synthesis genes (*pgcA*, *gtaB*, *ugtP*). We tested the growth and morphology of these strains and suppressor deletion/*mbl-KD* strains when possible. All strains except *tagO* and *tagD* were viable (Figure 5D). We quantified maximum growth rate and morphology of all strains using bulk growth measurements (area under the curve, AUC) and microscopy, respectively (Figures 5E, 5F, and 5G). As expected, the growth and morphological phenotypes of double-CRISPRi strains closely matched those of the equivalent suppressor deletion/*mbl-KD* strains (width *r*>0.90, *p*<10^-10^, AUC *r*>0.78, *p*<10^-6^; Figures 5E, 5F, and 5G). In general, the degree of growth and morphological rescue were correlated (Figure 5G). However, a few genes (*pgcA*, *ptsI*, and *glmR*) rescued only growth, consistent with a recent study showing that suppressing the lethality of *mbl* deletion does not necessarily require morphological compensation (Kawai et al., 2023). Taken together, these data suggest a role for *mbl* in WTA and LTA synthesis/attachment that impacts growth and cell-shape determination and is not shared with *mreB*.

### Identification of novel cell division genes

Cell division in bacteria is a highly orchestrated process in which constriction driven by the divisome machinery must be coordinated with cell-wall synthesis to avoid lysis (de Boer, 2010; Errington et al., 2003; Harry et al., 2006). To divide, cells form an FtsZ ring (Z-ring) at the site of the future septum that is used as a platform to assemble the divisome (Figure 6A), and the membrane constricts as PG is synthesized to form the septum and separate the daughter cells (Adams and Errington, 2009; Cameron and Margolin, 2024; Halbedel and Lewis, 2019). Since many cell division genes are essential, their GIs cannot be explored using a method dependent on knockouts. Our double-CRISPRi screen targeted essential genes with mismatched-sgRNAs that reduce but do not eliminate gene expression, enabling us to identify both positive and negative GIs of essential genes. Many divisome genes formed a highly connected network composed of strong negative GIs (Figures 6B and S7), consistent with the known co-dependence of these genes in cell division (Adams and Errington, 2009; Cameron and Margolin, 2024; Halbedel and Lewis, 2019). Moreover, we identified strong novel negative GIs between divisome genes and genes involved in PG precursor synthesis, PG remodeling, and TA synthesis/modification, as well as ECF sigma factors (SigX and SigM), reflecting the characterized activities of divisome proteins (Adams and Errington, 2009; Cameron and Margolin, 2024). By searching for additional genes that exhibited strong GIs or correlated GI profiles with known division genes, we also identified several potential new players in cell division, including the uncharacterized genes *yrrS*, *ytxG*, and *yerH* (Figure 5B).

**Figure 6.**
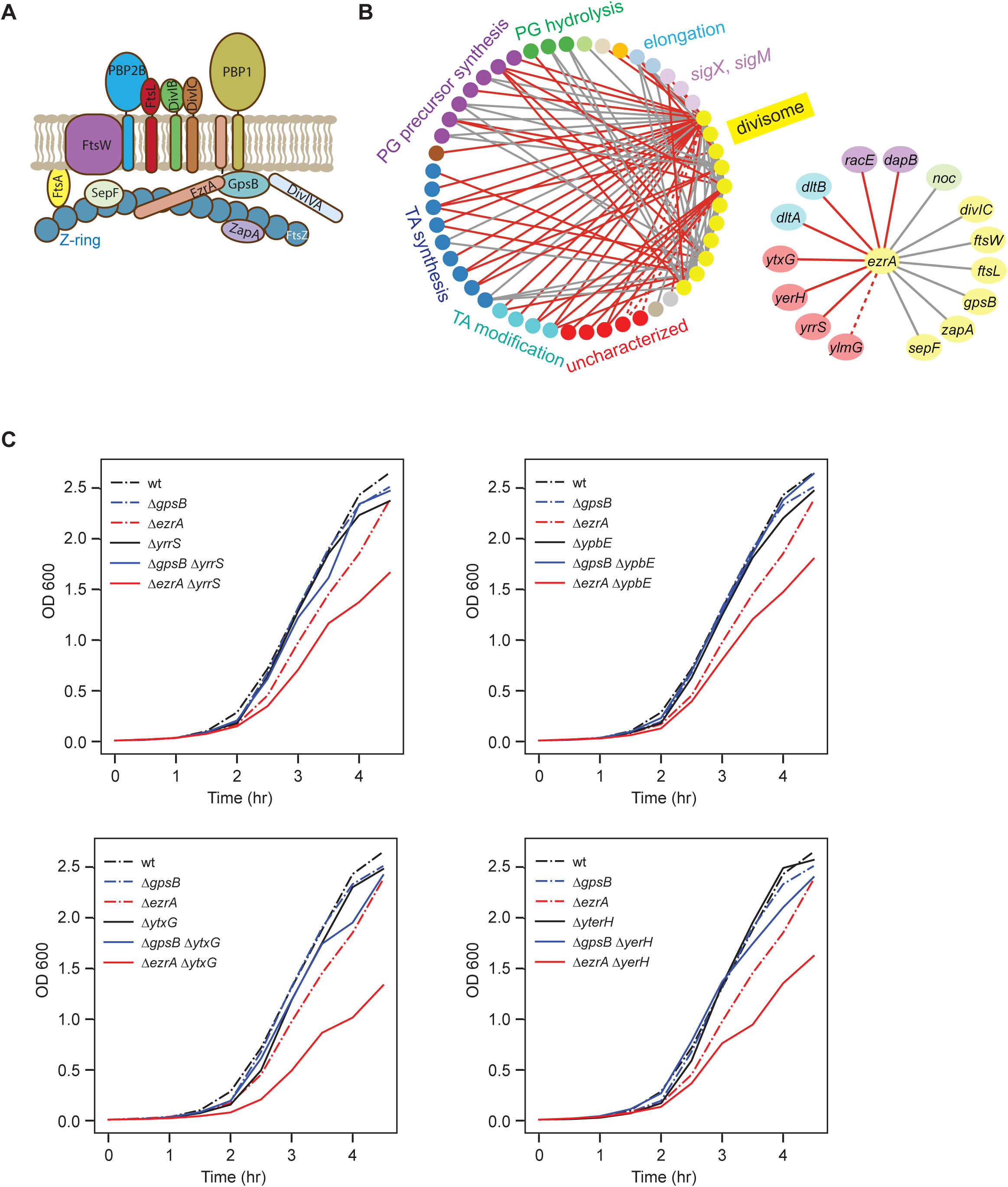
Double-CRISPRi identifies new players in *B. subtilis* cell division. **A)** Schematic of *B. subtilis* divisome complex showing the FtsZ ring, and associated cell wall synthesis complex (Halbedel and Lewis, 2019). **B)** Left: Schematic of intra- and inter-GI network of divisome genes. Right: GI network of *ezrA*. Gray lines indicate known interactions and red lines indicate novel interactions identified in this screen. The dotted line between *ezrA* and *ylmG* indicates false negative GI resulting from knockdown of *sepF* in the same operon due to the polar effect of CRISPRi. **C)** Deletion of novel cell division genes (*yrrS, ypbE, ytxG*, and *yerH*) exhibit synthetic growth phenotype with deletion of *ezrA*, but not with that of *gpsB*. Two independent experiments were performed and the representative data are shown here.

These uncharacterized genes (y-genes) all exhibited strong negative GIs with *ezrA*. EzrA is a negative regulator of Z-ring formation; in its absence, there are multiple Z-rings at the cell poles and mid-cell (Adams and Errington, 2009; Levin et al., 1999). EzrA also recruits PBP1 to the division septum (Claessen et al., 2008) and activates PrkC (Pompeo et al., 2015). We found that *ezrA* exhibited strong negative GIs with its known interaction partners *gpsB*, *sepF*, and *zapA*, whereas *yrrS*, *ytxG,* and *yerH* did not, raising the possibility that these uncharacterized genes function with one of the known *ezrA* interaction partners. Consistent with this hypothesis, YrrS in *B. subtilis* and YtxG in *S. aureus* have been reported to physically interact with GpsB (Bartlett et al., 2024; Cleverley et al., 2019). To validate these GIs, we constructed and characterized deletion strains of *yrrS*, *ytxG*, and *yerH* as well as *ypbE* (which was missing from our screen due to low sequencing read depth but has a similar protein-protein interaction profile to that of *yrrS* (Cleverley et al., 2019)) in a *ezrA* deletion strain. Using these double mutants, we found that all four y-genes exhibited negative GIs with *ezrA* but not *gpsB,* consistent with the results of our pooled screen (Figure 6C). Since YrrS and YpbE are known to bind together (Cleverley et al., 2019), we asked whether they interact synergistically with *ezrA*. Indeed, although the *yrrS*/*ypbE* double mutant exhibited no significant growth phenotypes, the *yrrS*/*ypbE*/*ezrA* triple deletion mutant was much sicker than predicted (Figures S8A and S8B). The interaction was specific to *ezrA*: *yrrS*/*ypbE* double mutants did not exhibit negative GIs with other *ezrA*-interacting cell division genes such as *gpsB*, *sepF*, and *zapA* (Figure S8B). These proteins may also have additional roles in division, as each had distinct but uncorrelated GIs (Supplementary Note 3, Table S4).

### Single-cell imaging reveals phenotypes for novel cell division gene knockouts

To further characterize the role of *yrrS*, *ypbE, ytxG*, and *yerH* in cell division, we imaged cells with deletions of these genes in exponential phase with or without CRISPRi knockdown of *ezrA* (Methods). In each case, we acquired phase-contrast, FM4-64, and DAPI images of thousands of cells and computationally segmented cell boundaries to capture cell, membrane, and nucleoid morphologies, respectively (Methods). Since YrrS, YpbE, and likely YtxG bind GpsB, we compared the morphologies of each strain to both *ezrA-KD* and *ezrA-KD*/Δ*gpsB* strains. As expected, *ezrA* knockdown resulted in increased filamentation as measured from phase-contrast images (∼32% increase in median cell length, Welch’s t-test *p* < 0.001, Figure 7A). However, FM4-64 membrane staining revealed that these apparently filamentous cells contained membrane cross-bands that demarcated compartments of length comparable to cells without *ezrA* knockdown (Figures 7A and 7B). Nucleoid localization, visualized with DAPI staining (Methods), was normal (Figure 7B). These data suggest that, in cells with *ezrA* knockdown, the cell division machinery assembles at the proper locations but is not fully functional.

**Figure 7:**
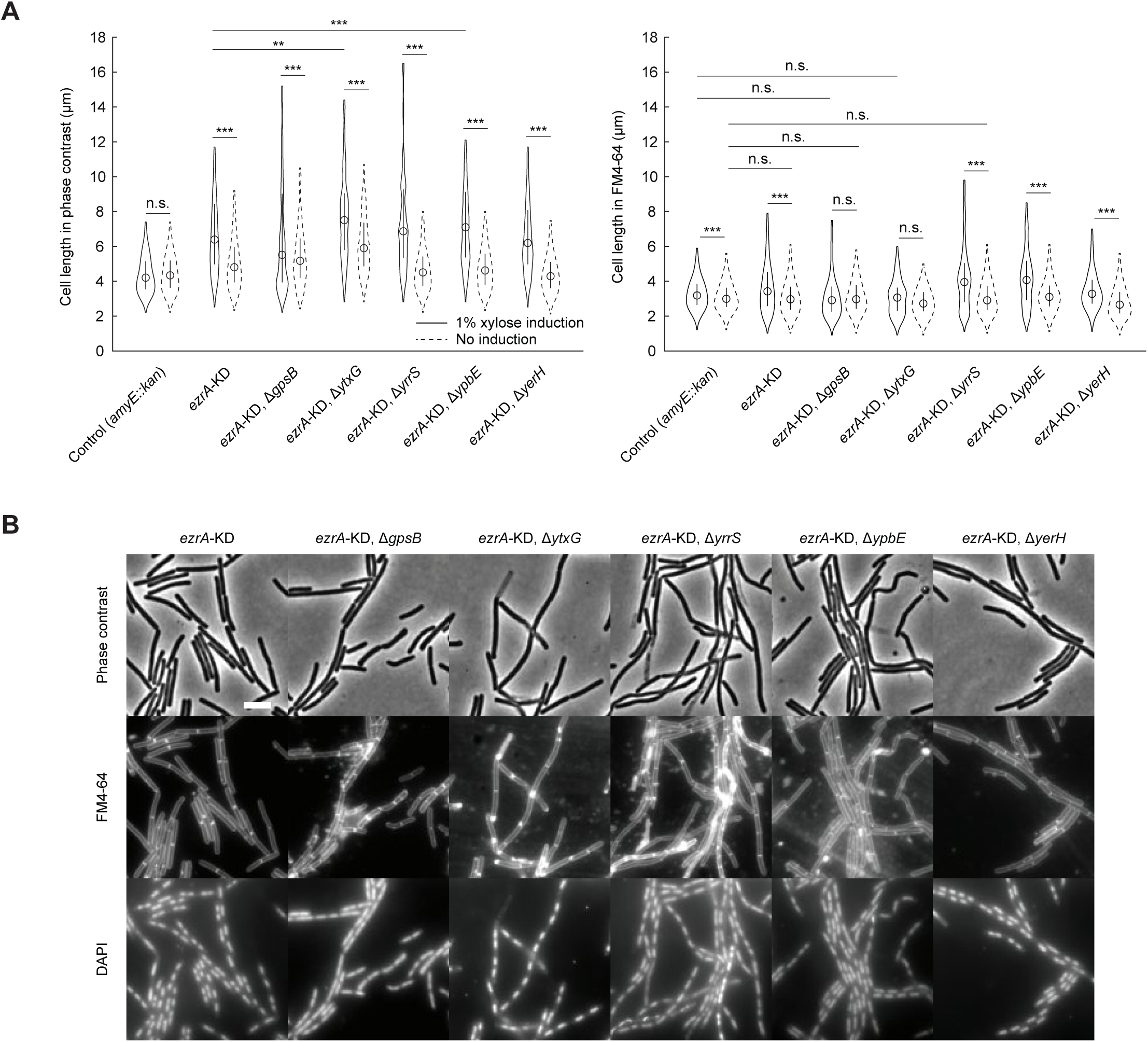
Knockout mutants of several uncharacterized genes exhibit defective cell pole synthesis under *ezrA* depletion. **A)** Depletion of EzrA causes cell elongation in Δ*gpsB* and several previously uncharacterized mutants (Δ*yerH*, Δ*ypbE*, Δ*yrrS*, and Δ*ytxG*), compared to *ezrA* depletion alone or gene knockouts without *ezrA* depletion, as indicated by phase-contrast microscopy. However, the length of cell compartments separated by FM4-64-labeled membranes showed limited increase compared to the control strain suggesting unperturbed Z-ring localization. **B)** The elongated mutant cells caused by *ezrA* depletion are separated into shorter compartments by division planes labeled with FM4-64. In all mutants, cell division planes are excluded from the DAPI-labeled nucleoid, as normally seen in wild-type. Scale bar: 5 µm.

The phenotypes of deletions of the four genes in an *ezrA-KD* strain were consistent with the GI data, and were indicative of varying defects. Deletion of either *gpsB* or *ytxG* alone resulted in filamentation (median cell length 19% and 37% longer than the control, respectively, Welch’s t-test *p* < 0.001, Figure 7A), and as previously reported (Bartlett et al., 2024), Δ*ytxG* exhibited numerous patches of membrane staining (Figure 7B). *ezrA* knockdown in the Δ*ytxG* background resulted in a substantial increase in median cell length (27%). In the Δ*gpsB* background, *ezrA* knockdown resulted in only a slight increase in median cell length (8%), consistent with previous work (Claessen et al., 2008) but the tail of the distribution extended to much longer lengths. Both strains, with or without *ezrA* knockdown, exhibited normal membrane cross-bands and nucleoid localization (Figure 7B). In contrast, the *yrrS* and *ypbE* deletions alone did not exhibit gross morphological defects, but upon *ezrA* knockdown exhibited increased filamentation relative to *ezrA* knockdown alone (∼8-12% longer median cell length, Welch’s t-test *p* < 0.001, Figure 7A). *ezrA* knockdown in these strains additionally resulted in increased distance between membrane crossbands (>20% larger than control, Welch’s t-test *p* < 0.001, Figure 7A), suggesting that these strains may have a mild but orthogonal defect in cell division that exacerbates the defects of *ezrA* knockdown. Finally, deletion of *yerH* alone lacked morphological phenotypes. Upon *ezrA* knockdown, this strain exhibited filamentation, membrane cross-bands, and nucleoid localization similar to that of the *ezrA* knockdown alone. However, *ezrA-KD*/Δ*yerH* strains exhibited increased cell wall bending and lysis compared to the *ezrA* alone (Figure 7B and Supplementary Note 4). Together, these data demonstrate the ability of GI analysis to reveal new genes involved even in well-studied processes like cell division, and highlight the diversity of phenotypes that can emerge from disruption of the division machinery.

## PERSPECTIVE

Here, we present double-CRISPRi, an experimental and analytical approach for high-throughput CRISPRi-based GI mapping in bacteria. We use double-CRISPRi to perform genome-scale GI-mapping of envelope-function genes, including essential genes, in the model bacterium *B. subtilis*. Our focus on mapping interactions between cell envelope-related genes allowed us to validate many of our findings using the vast existing knowledge base. This GI map serves as a broad resource for further characterization of envelope gene function, and our experimental and analytical framework will enable future GI mapping efforts in *B. subtilis* and other diverse bacteria. Our analysis of GIs in the *B. subtilis* cell-envelope supports three major conclusions.

First, we establish double-CRISPRi as a powerful tool for understanding bacterial gene function and pathway connections. The GIs of a gene accurately identify known and novel functional partners of the genes, enabling us to connect diverse processes and dissect complex pathways. This success is exemplified by our studies of *mbl* and *mreB*, which guide the elongasome. The elongation machinery contains a pair of synthetic lethal PG hydrolases, *lytE* and *cwlO*, which maintain the balance between peptidoglycan synthesis and disassembly that is essential for cell proliferation (Hashimoto et al., 2012). Although previous studies found that MreB and Mbl differentially activate these hydrolases (Dominguez-Cuevas et al., 2013), our study additionally uncovered an extensive network of GIs involving these genes. Moreover, our study identified extensive GIs between *mbl* (but not *mreB*) and many other processes, including LTA and WTA synthesis, the regulation of metabolism, and cell division that will motivate future studies. Our finding that *mbl* and other elongasome components genetically interact with division genes such as *divIVA, divIB, sepF, ftsL*, and *ftsA* is supported by a concurrent double-CRISPRi screen in *Streptococcus pneumoniae,* which found and validated negative GIs between *divIB*/*divIC* and many components of the elongasome (Dénéréaz et al., 2024 co-submitted).

Second, we establish the ability to identify new members of essential cellular machines. Our screen leveraged mismatch-CRISPRi (Hawkins et al., 2020) to design sgRNAs that target essential genes with intermediate efficacy, resulting in single mutants with quantifiable growth rates that enabled the identification of both positive and negative GIs. A striking example of this was the identification of additional players in the well-characterized and intensively studied process of cell division. Cell division genes, including many essential genes such as *divIB* and *ftsL*, formed a highly interconnected network of GIs. We identified and characterized four genes connected to this cluster, highlighting the utility of GI mapping for discovering the complete network of divisome interactions. Single-cell imaging of these mutant strains revealed division defects not caused by the inhibition of septum formation, with morphological defects suggesting overstabilization of the division machinery and mis-localization of growth at sites of intended septa, as has been observed in *ezrA gpsB* double mutants (Claessen et al., 2008) (Supplementary Note 4). Our GI data support a role for these genes in cell division via localization of PBP1 (Supplementary Note 3). Three of these genes (*yrrS*, *ypbE*, and *yerH*) are conserved primarily in *Bacillus* species and closely related genera, suggesting a specialized function in the division machinery of these species. However, *ytxG* is broadly conserved in both rod-shaped and coccoid Firmicutes, and exhibits distinct phenotypes in each. Together, these data suggest that while the core cell division machinery is highly conserved (Adams and Errington, 2009), accessory factors and PPIs can differ across taxa. Future double-CRISPRi studies in diverse bacteria will reveal how the cell division machinery has been adapted to different cell shapes (rod, cocci, spiral), modes of cell wall growth (symmetric division, apical growth), and bacterial lifestyles.

Third, at a broader level, our screen begins to reveal the nature and frequency of GI in bacteria, which informs and constrains future studies. As expected based on GI studies in yeast (Costanzo et al., 2016), essential and well-characterized genes (e.g. gene set 1; ∼3.8 GIs with |GI score| > 2) exhibited more GIs than uncharacterized genes (e.g. gene set 2; ∼0.5 GIs with |GI score| > 2), suggesting they may function as network hubs. This highlights the utility of targeting essential genes with mismatched sgRNAs and ensuring high library coverage and sequence depth to accurately quantify strong growth defects. Moreover, the surprising number of inter-process connections argues for selecting broad gene sets rather than focusing on single processes in future studies.

Our study provides a valuable data set for deciphering cell envelope gene function in *B. subtilis* and a blueprint for studies in other bacteria. The double-CRISPRi approach is a robust tool for deeper and broader study of bacterial GI networks, which can illuminate new biology and enable rational design of antibiotic combination therapies. Double-CRISPRi libraries can be used to (1) conduct chemical genomic screens that reveal multi-partite interactions and phenotypes for highly redundant genes, (2) can be combined with mobile-CRISPRi (Peters et al., 2019) to study GIs in diverse bacteria, and (3) could (with modifications, see Limitations) be scaled to target all genes in a bacterial genome. Additionally, our current envelope-focused double-CRISPRi library can be combined with high-throughput microscopy or flow-cytometry approaches (Bartlett et al., 2024; Juillot et al., 2021; Shi et al., 2017) to assay cell shape, size, and other, non-growth-related phenotypes. Double-CRISPRi will serve as an important tool for closing the gene sequence-function gap across bacterial species.

### Limitations

The substantial insights into cellular connectivity enabled by our double-CRISPRi method motivate future efforts to target the entire genome. In our study, we individually cloned the first sgRNA to ensure even representation in the double mutant pool.

However, genome-wide targeting requires pooled cloning of sgRNAs at both positions, which can be accomplished using optimized plasmids (pDCi00, Table S5). Importantly, targeting every pairwise combination of the ∼4,000 genes (∼16 million strains) in a typical bacterial genome would require growth of large-volume cultures to avoid bottlenecking and would entail proportionately higher sequencing costs. To mitigate these issues, a double-CRISPRi library could be designed to target only the first gene in an operon, relying on CRISPRi polarity to repress downstream genes (Peters et al., 2016). As ∼50% of bacterial genes are in operons (Geissler et al., 2021), such a strategy would reduce library size ∼4-fold. However, only computational predictions of operon structure are available for many species. These predictions incorrectly annotate some operon boundaries and can miss (conditional) internal promoters. Indeed, the discordant GIs of operon members *pbpI* and *yrrS* are likely due to a promoter upstream of *yrrS* unaffected by *pbpI* knockdown. Therefore, it is likely prudent to target each gene individually, which should be increasingly tractable as advances in sequencing and synthesis technology reduce the associated costs.

## METHODS

### Strains and growth conditions

All strains used in this study are listed in Table S5. All *B. subtilis* strains were derivatives of the 168 strain (Bacillus Genetic Stock Center; accession number: 1A1). Cells were routinely grown in lysogeny broth (LB) medium (1% tryptone, 0.5% yeast extract and 0.5% NaCl) at 37 **°**C with aeration or on LB agar plates supplemented with appropriate antibiotics at the specified concentrations (by activity) if needed: For *B. subtilis*, erythromycin (1μg/ml), lincomycin (12.5μg/ml), spectinomycin (100μg/ml), chloramphenicol (6μg/ml), kanamycin (7.5μg/ml). For *E. coli*, carbenicillin (100μg/ml).

### Genetic manipulation

Transformation of the plasmid into *E. coli* strain was performed using the heat shock method or electroporation as described in the New England Biolabs (NEB) protocol (https://www.neb.com/en-us/protocols/0001/01/01/high-efficiency-transformation-protocol-c3019, https://www.neb.com/en-us/protocols/0001/01/01/electroporation-protocol-c3020).

Transformation of *B. subtilis* was performed using natural competence. Competent cells were prepared by following protocol (Koo et al., 2017); *B. subtilis* cells were inoculated into 3 ml of MC medium (10.7 g/L K_2_HPO_4_, 5.2 g/L KH_2_PO_4_, 20 g/L glucose, 0.88 g/L trisodium citrate dihydrate, 0.022 g/L ferric ammonium citrate, 1 g/L casamino acids, 2.2 g/L potassium glutamate monohydrate, 20 mM MgSO_4_, 300 nM MnCl_2_, 20 mg/L L-tryptophan) and incubated at 37 **°**C overnight with aeration. The overnight culture was diluted to an OD_600_ of 0.1 in 20 ml competence medium (10.7 g/L K_2_HPO_4_, 5.2 g/L KH_2_PO_4_, 20 g/L glucose, 0.88 g/L trisodium citrate dihydrate, 0.022 g/L ferric ammonium citrate, 2.5 g/L potassium aspartate, 10 mM MgSO_4_, 150 nM MnCl_2_, 40 mg/l L-tryptophan, 0.05% yeast extract), then grown in a 125 ml flask at 37**°**C with shaking (250 rpm) until cells reached OD_600_∼1.5. 120 µl of culture was then mixed with up to 10 µl DNA and incubated at 37 **°**C with shaking. After 2 hr of incubation, cells were plated on LB agar containing selective antibiotics.

When needed, the kanamycin resistance cassette flanked by *lox* sequences was removed using Cre recombinase as previously described (Koo et al., 2017). Briefly, a strain containing the *lox* flanked kanamycin resistance cassette was transformed with pDR244 (a temperature-sensitive plasmid with constitutively expressed Cre recombinase). Transformants were selected on LB agar plates supplemented with 100 μg/mL spectinomycin at 30 °C. Transformants were then streaked on LB agar plates and incubated at 45 °C. Cells from the edge of single colonies were then restreaked on LB, LB supplemented with kanamycin, and LB supplemented with spectinomycin.

Strains that grew on LB agar plates, but not on LB agar plates supplemented with antibiotics, had lost pDR244 and the *lox*-flanked kanamycin resistance cassette.

### Construction of a new dcas9 expressing strain

BKC30001 was constructed by replacing the erythromycin-resistance gene of our previously described *dCas9* strain CAG74209 (Peters et al., 2016) with a fragment containing kanamycin-resistance cassette flanked with *lox* sites that was generated by joining three PCR fragments: the kanamycin resistance cassette, and 1kb 5’ and 3’ flanking regions of the erythromycin-resistance gene in CAG74209. The kanamycin resistance cassette in pDR240a was amplified using primers oDCi005 and oDCi006. 1kb 5’ and 3’ flanking regions of the erythromycin-resistance gene in CAG74209 were amplified using the oDCi001/0DCi002 primer pair and the oDCi003/oDCi004 primer pair respectively. Amplified DNA fragments were purified using Agencourt AMPure XP (Beckman Coulter, Cat# A63881) magnetic beads. The purified DNA fragments were mixed and subjected to the joining PCR using oDCi003 and oDCI006. The joined PCR product was transformed into CAG74209. Three kanamycin-resistant but erythromycin-sensitive clones were isolated and their genomic DNA was purified using the Qiagen DNeasy Blood & Tissue kit (Cat# 69506). The sequence of *dcas9* was verified by Sanger sequencing using primers (Table S6). The confirmed genomic DNA was re-transformed into the wild-type 168 strain, generating BKC30001.

### Construction of double sgRNA plasmids

The double sgRNA plasmid pBsuDCi was modified from pDG1662. The pool of pBsuDCi was constructed through three major steps.

First, to increase transformation and double-crossover efficiency, 1.5kb of DNA upstream of *amyE* was PCR amplified from *B. subtilis* 168 genomic DNA and inserted into pDG1662 via HiFi Assembly (all enzymes and reaction kits used in cloning were purchased from NEB, and high-fidelity versions of restriction enzymes were used if available), replacing the shorter upstream fragment of *amyE* in pDG1662. The synthetic DNA (IDT) containing a transcription terminator, P*veg* with BbsI and P*scr* with BsaI cut sites for spacer cloning, random barcode sequence, and downstream tandem transcription terminators was cloned into the previously described pDG1662 derivative via HiFi Assembly. The annealed oligonucleotide containing sgRNA sequence targeting *yabE* with flanking restriction sites was ligated with the purified plasmid digested with BbsI, generating pBsuSCi0.

Second, using pBsuSCi0 as a template, the fragments containing sgRNA1(Figure 1A and Table S1) and associated random barcodes were individually generated by PCR using the primer pairs of a sgRNA-specific oligonucleotide, oDCi_sgRNA1 (5’ TGTACAATAAATGT-sgRNA sequence-GTTTTAGAGCTAGAAATAGCAAGTTA 3’) and a random barcode containing oligonucleotide, oDCi014 (5’ GGCGCGGCCGCAAAACAAGAAAGAGAAAAGTTCCCTATNNNNNNNNNNNNNNNNNNNNNNNNNNAGATCGGAAGAGCACACGTC 3’). Each purified fragment was digested with BsrGI and EagI, and cloned into pBsuSCi0 which was digested with the same enzyme followed by dephosphorylation, individually generating a series of pBsuSCi containing sgRNA1 and barcode. Barcodes associated with each sgRNA1 were then identified via Sanger sequencing of purified plasmids. The purified equimolar plasmids were pooled in 7 tubes, each of which contained 45∼50 sgRNAs.

Finally, sgRNA2s (Figure 1A and Table S1) were cloned into the BsaI sites of pooled double sgRNA plasmids (pBsuSCi) that contained cloned sgRNA1s. sgRNA2 fragments were prepared in two ways. One fraction of sgRNAs was prepared by individually annealing two single-stranded DNA oligonucleotides to create 4-base overhangs, followed by pooling. The rest of the sgRNAs were prepared via digestion of pooled sgRNA fragments with BsaI. To generate sgRNA fragment pools, oligonucleotide pools containing the sgRNA spacers with flanking restriction sites and PCR adapters were obtained from Agilent Technologies. The oligonucleotide pools were amplified via 14 cycles of PCR using Q5 DNA polymerase and primers. The purified PCR product was digested with BsaI-HFv2 and purified after PAGE in 10% TBE gels (Invitrogen Cat# EC6275BOX) to remove adapter ends. Seven pBsuSCi plasmid pools were individually digested with BsaI-HFv2 for 2 hr. Final double sgRNA plasmid libraries were constructed in two ways depending on the inserts. For the inserts prepared by annealing, the equimolar digested vector pools were combined and ligated with the inserts. In contrast, for the inserts prepared by digestion with BsaI, each vector pool was dephosphorylated and ligated with inserts individually. Each ligation was carried out using 100 ng of digested vector at a 1:2 (vector: insert) molar ratio for 3 hr at 16 **°**C using T4 DNA ligase. Each of the 8 ligated products was transformed into electrocompetent cells (NEB #C3019), and cells were recovered in SOC medium at 37°C for 1 hr, then inoculated into 100 ml of LB with carbenicillin and grown overnight. Each plasmid library was purified using a midiprep kit (Qiagen, Cat# 12143).

### Construction of the *B. subtilis* double-CRISPRi library

The double CRISPRi library was constructed by transforming double sgRNA plasmid libraries into BKC30001 using natural competence. The 8 pools of plasmids were linearized via NdeI digestion before transformation to eliminate single-crossover recombination. To increase the transformation scale, the protocol was modified as follows. 300 ng of digested plasmid DNA were mixed with 120 µl of fresh competent cells and incubated in deep 96-well plates. After incubation at 37 °C for 2 hr with shaking (900 rpm), 10 reactions were combined in Eppendorf tubes, and cells were spun down at 5000 *g* for 1 min. After discarding 900 µl of supernatant from each tube and resuspending cells, cells were plated on LB agar plates supplemented with chloramphenicol to select for plasmid integration, and the plates were incubated at 37°C for 16 hr. The yield of each batch of transformation was calculated from CFU counting after serial dilution. The average plating density was ∼0.3 X10^6^ CFUs/plate and the total number of transformants was more than 100 times the library size. To store the library, plates were scraped, pelleted, and resuspended in S7 salts with 12.5% glycerol, and stored in 500 µl aliquots at -80 °C. The number of clones in each aliquot was calculated by measuring the OD_600_ of the aliquot after serial dilution.

### Fitness experiments and preparation of Illumina sequencing libraries

Growth experiments with the *B. subtilis* double-CRISPRi library were performed in triplicate and samples were taken as described in Figure S2. Glycerol stocks of 8 pools of the library were fully thawed and inoculated into 500 mL of LB at an OD_600_ of 0.04.

These cultures were grown to an OD_600_ of 0.32 at which point all cultures were combined to evenly distribute all clones in one tube. This culture was set as the T0 time point sample. The T0 culture was diluted to an OD_600_ of 0.01 in1 liter of fresh LB + 1% xylose and then repeatedly grown to an OD_600_ of 0.32 (∼5 doublings) followed by dilution to an OD_600_ of 0.01 a total of 3 times (to enable 15 doublings), resulting in samples T1, T2, and T3. For the overnight growth and recovery screen, the T0 culture was diluted to an OD_600_ of 0.01 in 1 liter of LB (samples T4 and T5) or LB + 1% xylose (samples T6 and T7) and then grown for 18 hr. Each overnight culture (T4 and T6) was diluted to an OD_600_ of 0.01 in 1 liter of fresh LB + 1% xylose and then grown to an OD_600_ of 0.32 (∼5 doublings, samples T5 and T7). 1 ml of the culture volume was collected immediately before dilution and after the final growth phase. Cells were pelleted by spinning down at 15000*g* for 2 min in Eppendorf tubes and stored at -80 °C.

Genomic DNA of the cell pellets was purified using a Qiagen DNeasy Blood & Tissue kit. The sequencing region was amplified from 2 µg of genomic DNA (1000X coverage of each clone) using Q5 DNA polymerase for 14 cycles with primers harboring distinct indices for different replicates and sampling times (Table S6). Differentially indexed PCR products were purified after PAGE in 8% TBE gels and combined at an equimolar ratio. The combined sample was split into three lanes for sequencing on a Novaseq 6000 with 100 bp paired-end reads at the UCSF Center for Advanced Technology using custom sequencing primers (Table S6).

### Relative fitness (RF) quantification

Raw FASTQ files were aligned to the library oligos and enumerated using the script at https://github.com/traeki/mismatch_crispri, count_guide_pairs_2021.py, pseudocounts of 1 were added, and relative fitness was calculated as previously described (Hawkins et al., 2020). Briefly, for each strain *x* with at least 100 counts at *t*_0_, we calculate the relative fitness *F*(*x*) according to

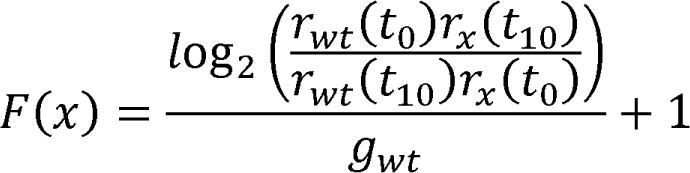

where *r_x_*(*t_i_*) is the fraction of strain *x* in the population at time *i* and *g_wt_* is the number of generations of wildtype growth in the experiment. In our experiments, *g_wt_* was calculated from the OD measurements of the culture, and *r_wt_*(*t_i_*) was calculated as the median of 2024 strains with non-targeting sgRNAs at both positions.

### GI score calculation and filtering

#### Calculating expected fitness

We used an additive model to calculate expected fitness. The fitness defects (1-RF) of each parent strain were added together and subtracted from 1.

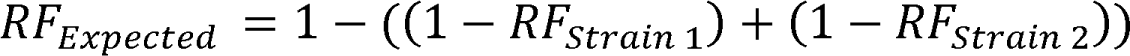

An additive model was chosen over a multiplicative model (Mani et al., 2008) for two reasons. First, an additive model makes reasonable predictions if one or both parent strains has negative RF. For example, if RF_parent A_ = -0.5 (i.e., the strain is diluted from the pool faster than dilution, for example via lysis) and RF_parent B_ = 0.5, a multiplicative model would give an expected RF for the double mutant of -0.25, which is less sick than parent A, an illogical conclusion. The situation is even worse if both parent strains have a negative fitness: the expected fitness would then be positive. Second, for the most frequently encountered fitness defects (1 > RF > 0.75), an additive and multiplicative model give similar results. Consider RF_parent A_ = 0.9 and RF_parent B_ = 0.9. The additive definition predicts an expected RF for the double mutant of 0.8, while the multiplicative definition gives 0.81. Our choice is supported by the literature (Mani et al., 2008) and by a concurrent study (Dénéréaz et al., 2024 co-submitted).

#### GI score calculation

We used a custom R code (https://github.com/horiatodor/GI-Score) to determine GI scores. A schematic We first identified a set of control strains. To do so, we considered the median across all rows (sgRNA1) and all columns (sgRNA2). “Control” columns were those with column medians within 1 MAD of the median of column medians, and “Control” rows were those with row medians within 1 MAD of the median of row medians. For each double-CRISPRi strain we then calculated a distribution of expected RF by adding the fitness defect of all “control” rows and all “control” columns. The GI score was then calculated as the robust (median, MAD) *z*-score of the measured strain fitness. GI scores were calculated separately for the 316×333 library and the 316×982 library, since these were constructed separately. The GI scores were independently calculated for each of three biological replicates and averaged.

#### Filtering

During the course of our analysis, we found that several sgRNAs had many GIs and that these GIs were correlated with each other. Since these GIs appeared to be due to a systemic artifact, we searched for a technical explanation that would allow us to filter these sgRNAs from the dataset. We found that the GIs of these sgRNAs were highly correlated to the PAM-distal sequence of the interacting sgRNA, which is suggestive of an issue with sgRNA transcription (these bases serve as the transcription start site). We eliminated these spurious hits as follows. For each sgRNA in position 1 (rows), we performed a linear regression between its GI scores and the one-hot encoded first 2 nucleotides of the interacting sgRNA. We constructed a distribution of correlations, and eliminated all sgRNAs with a correlation greater than the median plus 5 times the MAD of the distribution of correlations. The same process was applied to sgRNAs at position 2 (columns). Approximately 10% of strains were filtered through this process.

#### Correlation matrix calculation

The correlation of GI scores was calculated as the Pearson correlation between all sgRNAs at position 2 (columns), resulting in a matrix of 1315 × 1315 correlations.

#### STRING analysis

Interactions from the STRING database were retrieved from https://string-db.org, for *Bacillus subtilis* strain *168* (taxid: 224308).

### High-throughput imaging

Cells from frozen stocks were diluted 1:30 into 300 µl of LB in a deep 96-well plate (Beckman Coulter, #267007), covered with a breathable film, and incubated at 37 °C with shaking at 1000 rpm. After 3 hr of incubation, the culture was diluted to OD_600_∼0.01 into LB with 1% xylose to induce knockdown of target gene(s) or without xylose and further incubated in a 96-well flat-bottom plate (Greiner Bio-One, #655180) at 37 °C with shaking at 1000 rpm. After 3 hr of incubation, the culture was passaged again in LB with or without 1% xylose and further incubated in a 96-well flat-bottom plate at 37 °C with shaking at 1000 rpm. OD_600_ was measured using a Biotek Epoch plate reader to monitor growth during the two passages after the initial inoculation. Cells were then transferred from 96-well plates to 1% agar pads with 0.85X PBS using a 96-pin array (Singer Instruments, Cat# REP-001) and imaged using SLIP, a previously described high-throughput single-cell imaging protocol (Shi et al., 2017). Phase-contrast images were acquired with a Ti-E inverted microscope (Nikon Instruments) using a 100X (NA 1.40) oil immersion objective and a Neo 5.5 sCMOS camera (Andor Technology). Images were acquired using μManager v. 1.4 (Edelstein et al., 2010).

### Cell staining and imaging

After growing the cells in LB with or without induction, they were transferred to an LB agarose pad containing 1% agarose. FM4-64 and/or DAPI were added directly to the agarose pad at final concentrations of 5 µg/mL and 1 µg/mL, respectively. The cells were then imaged using a Nikon Ti-E inverted microscope equipped with a 100X (NA 1.40) oil immersion objective and a Prime BSI Express sCMOS camera (Teledyne Photometrics).

### Microscopy image analysis

Phase-contrast and fluorescence images were analyzed using the MatLab software *Morphometrics* (Ursell et al., 2017). A local mesh grid was generated for each cell contour using a method adapted from *MicrobeTracker* (Sliusarenko et al., 2011) to obtain cell length and width. For each cell, length was determined as the distance along the centerline between the two poles. Lysed cells and cells with fluorescent foci or specific shape defects were manually counted to estimate the frequency of lysis/shape defect in certain mutants.

### Whole-genome sequencing of secondary suppressor strains

Secondary suppressor strains were obtained by transformation of a *mbl*::*kan* fragment into suppressor deletion strains. Many *mbl*-suppressor double-deletion strains, as well as the triple-deletion strains harboring Δ*sigI*, lysed after overnight growth. Secondary suppressor strains regrew from lysed colonies (Supplementary Note 1), and were purified by picking cells from the regrown colonies followed by restreaking on a fresh LB agar plate. Purified single colonies were grown to an OD_600_ of 1 in LB. 1 ml of each culture was pelleted by spinning down at 15000*g* for 2 min in Eppendorf tubes. Genomic DNA of the cell pellets was purified using the Qiagen DNeasy Blood & Tissue kit. Purified DNA was submitted to Seqcenter (Pittsburgh, PA, USA) for sequencing with Illumina 2×151 paired-end sequencing and identifying the mutations.

## ACKNOWLEDGMENTS

We thank Garner E.G., Veening J.-W., Vollmer W., and members of Gross laboratory for their helpful comments. This study was supported by NIH R35GM118061 to C.A.G., NIH R35GM150487 to J.M.P., Bio-X Stanford Interdisciplinary Graduate Fellowship to J.S., NIH RM1GM135102 to K.C.H., NIH R01AI147023 to K.C.H., and NSF EF-2125383 to K.C.H.. K.C.H. is a Chan Zuckerberg Biohub Investigator.

## AUTHOR CONTRIBUTIONS

Conceptualization, B.M.K., H.T., K.C.H., J.M.P., and C.A.G.; Methodology, B.M.K., H.T., K.C.H., J.M.P., and C.A.G.; Investigation, B.M.K., J.S.,J.S.H., C.C.H., A.B.B, and J.M.P.; Formal Analysis, B.M.K., H.T.,J.S.,J.v.G., K.C.H., J.M.P., and C.A.G, and C.A.G.; Writing, B.M.K., H.T.,J.S.,J.v.G., K.C.H., J.M.P., and C.A.G.; Funding Acquisition, K.C.H., J.M.P., and C.A.G.; Supervision, B.M.K., H.T., K.C.H., J.M.P., and C.A.G.

## SUPPLEMENTAL INFORMATION

Documents S1. Supplemental Figures S1 to S8

Documents S2. Supplementary Notes 1 to 4

Table S1. List of target genes, sgRNA sequences, and their associated barcode sequences, related to Figure 1

Table S2. List of RFs of the double-CRISPRi strains, related to Figures 1, 5, S3, and S4

Table S3. List of raw and filtered GI scores of the double-CRISPRi strains, related to Figure 2, 3, 4, 5, 6, S3, S4, S5, and S7

Table S4. List of GI correlations between genes (sgRNAs), related to Figure 3

Table S5. List of strains and plasmids used in this study, related to Figures 5, 6, 7, S6, S8, and Methods

Table S6. List of primers used in this study, related to Figure 1 and Methods

