## Supplemental Figures for "Comprehensive double-mutant analysis of the *Bacillus subtilis* envelope using double-CRISPRi"

**Supplementary Figures**

**Figure S1.** Related to Figure 1. **Chromosomally-integrated double-CRISPRi system in *B. subtilis.*** A) Detailed nucleotide sequence of sgRNA locus. Each genetic feature is indicated under sequence. B) Transcriptional profiles of *veg* and *scr* of which promoters were used for transcription of sgRNA1 and sgRNA2 respectively. Data was obtained from SubtiWiki (Pedreira et al., 2022). C) Distribution of read counts of sgRNA pairs in the three replicates of T0 samples (Fig S2). Distribution is presented by the box plot. Red line indicates the median of the read counts (replicate 1; 592, replicate 2; 546, replicate 3; 469)

**Figure S2**. Related to Figure 1. **Schematic of the pooled growth experiment and sampling time points.** Blue arrows indicate growth without xylose, and red arrows do growth with 1% xylose for CRISPRi induction. The replicates were obtained from the cultures of three independent culture flasks. Each culture size was 1 liter.

**Figure S3.** Related to Figure 1. **The RFs of the strains obtained in this study are highly reproducible.** A) Correlation between RF of three replicates. B) Correlation between RF of single gene knockdown strains in this study and the relevant strains in previously published single-CRISPRi experiments (Hawkins et al., 2020). Blue and orange dots indicate knockdown induced by sgRNA under P*veg* and P*scr* respectively.

**Figure S4.** Related to Figure 2. **Calculation of GI scores**. See Methods.

**Figure S5.** Related to Figure 2. **Double-CRISPRi identifies GIs.** A) Correlation between all GI scores of three replicates. Note that correlations are higher with a high GI score threshold. (GI score$r$ ~ 0.79 for at least one replicate with |GI-score|>3; $r$ ~ 0.59 for at least one replicate with |GI-score|>2). B) Distribution of the GI correlations between the genes within the same operon. C) GI scores for all strains with *mbl* (x-axis) and *mreB* (y-axis).

**Figure S6.** Related to Figure 4. **The defect in D-alanylation significantly impairs the growth of Δ*ltaS* strain but not that of Δ*yfnI* strain.** *dlt* operon deletion (*dltABCDE*::kan) and *amyE::kan* fragments (as a transformation control) were transformed into antibiotic marker-free Δ*ltaS* strain and Δ*yfnI* and incubated for 16 hours. Colonies of Δ*ltaS* Δ*dltABCDE* double mutant were not visible on the plate after 16 hours of incubation but were visible after 40 hours.

**Figure S7.** Related to Figure 6. **A detailed GI network of divisome genes.** The network is identical to Figure 6B. Gray lines indicate known interactions and red lines indicate novel interactions identified in this screen.

**Figure S8.** Related to Figure 6. **Deletion of *yrrS* and *ypbE* exhibit synthetic growth phenotype with deletion of *ezrA*, but not with those of other *ezrA*-interacting cell division genes such as *gpsB*, *sepF*, and *zapA*. A)** Growth phenotypes of Δ*yrrS* Δ*ezrA* and Δ*ypbE* Δ*ezrA*. **B)** Growth phenotypes of triple mutants. Colonies of Δ*yrrS* Δ*ypbE* Δ*ezrA* triple mutant were not visible on the plate after 16 hours of incubation but were visible after 40 hours.

### Figure S1

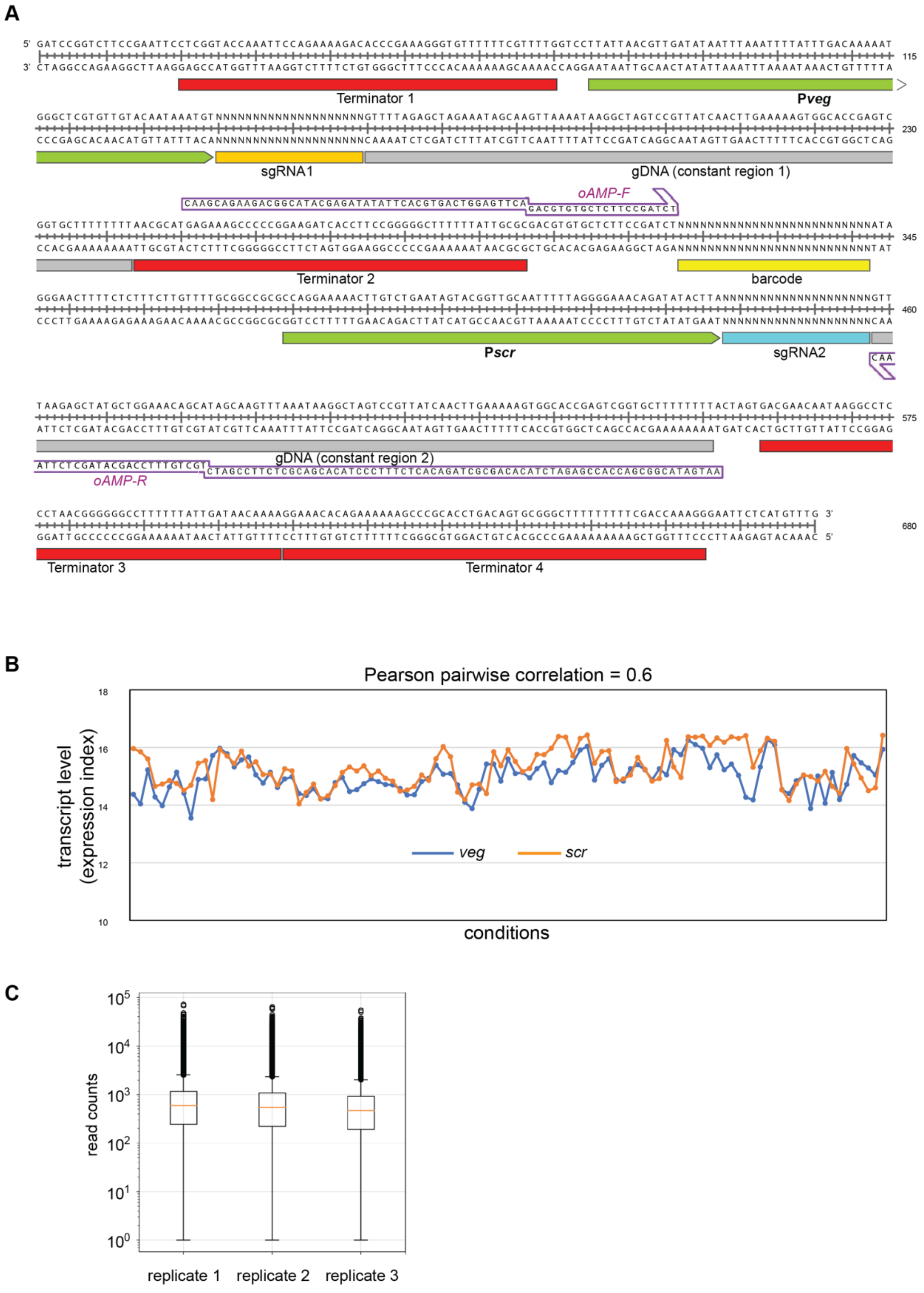

A
Terminator 1
Pveg
sgRNA1
gDNA (constant region 1)
oAMP-F
Terminator 2
barcode
Pscr
sgRNA2
gDNA (constant region 2)
oAMP-R
Terminator 3
Terminator 4
B
Pearson pairwise correlation = 0.6
transcript level (expression index)
conditions
veg
scr
C
read counts
replicate 1
replicate 2
replicate 3

**Figure S2**

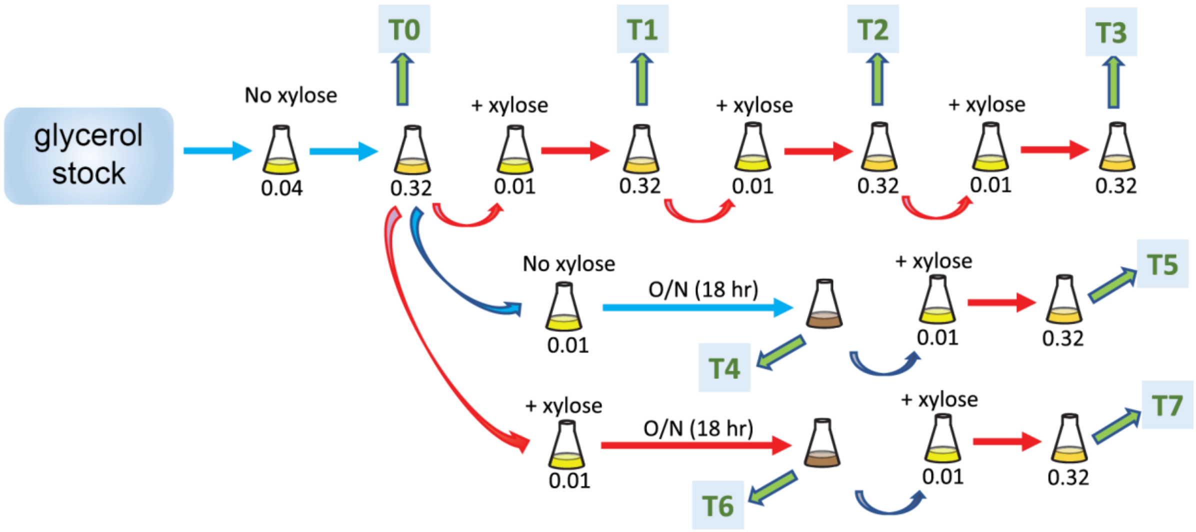

glycerol stock
No xylose
0.04
T0
0.32
+ xylose
0.01
T1
0.32
+ xylose
0.01
T2
0.32
+ xylose
0.01
T3
0.32
No xylose
0.01
O/N (18 hr)
T4
+ xylose
0.01
0.32
T5
+ xylose
0.01
O/N (18 hr)
T6
+ xylose
0.01
0.32
T7

**Figure S3**

**A**

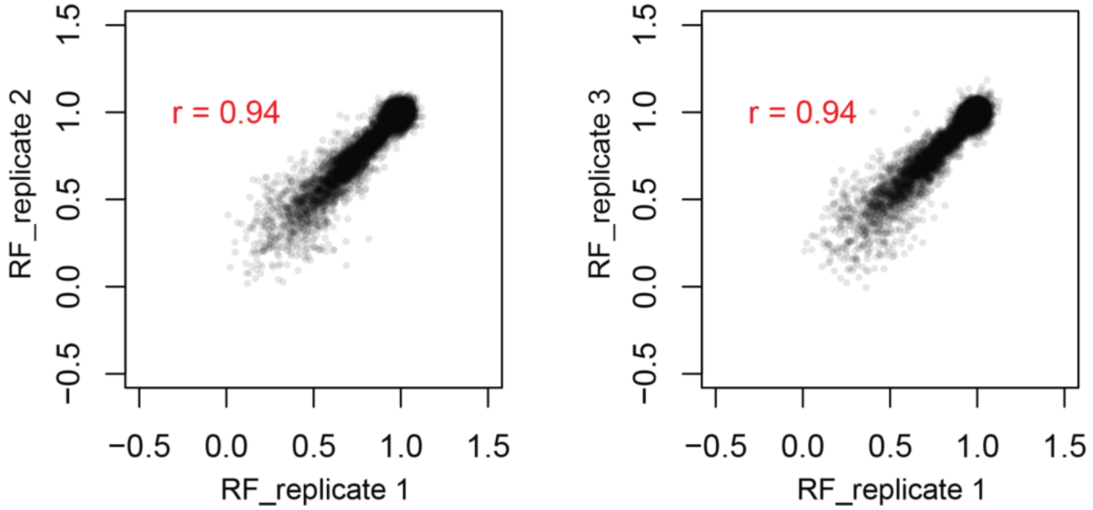
r = 0.94
RF_replicate 2
RF_replicate 1
r = 0.94
RF_replicate 3
RF_replicate 1
−0.5 0.0 0.5 1.0 1.5

**B**

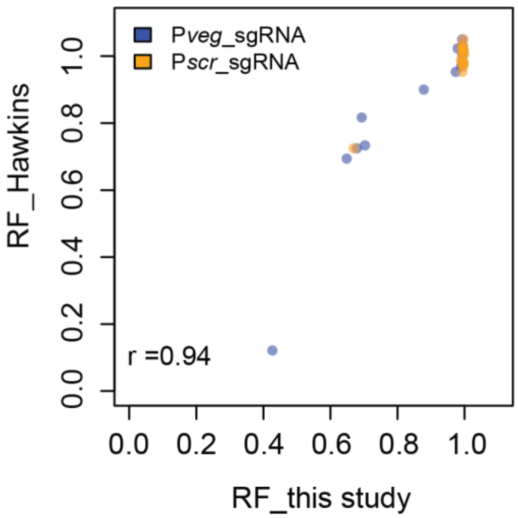
Pveg_sgRNA
Pscr_sgRNA
r =0.94
RF_Hawkins
RF_this study
0.0 0.2 0.4 0.6 0.8 1.0

**Figure S4**

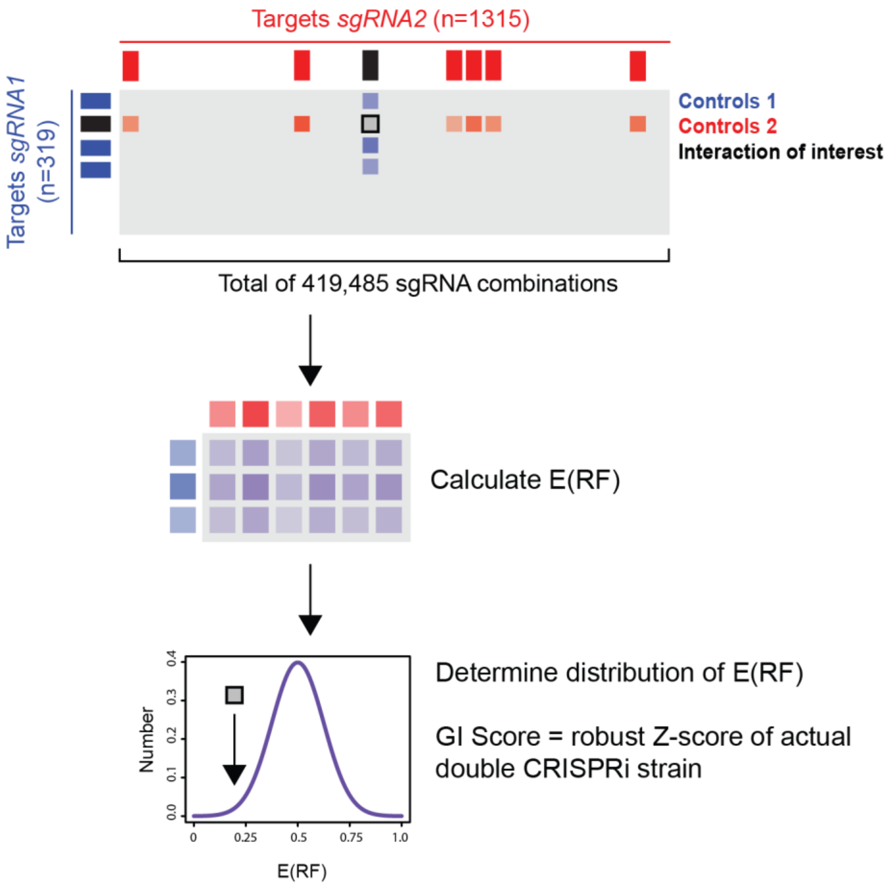


Determine distribution of E(RF)

GI Score = robust Z-score of actual double CRISPRi strain

**Figure S5**

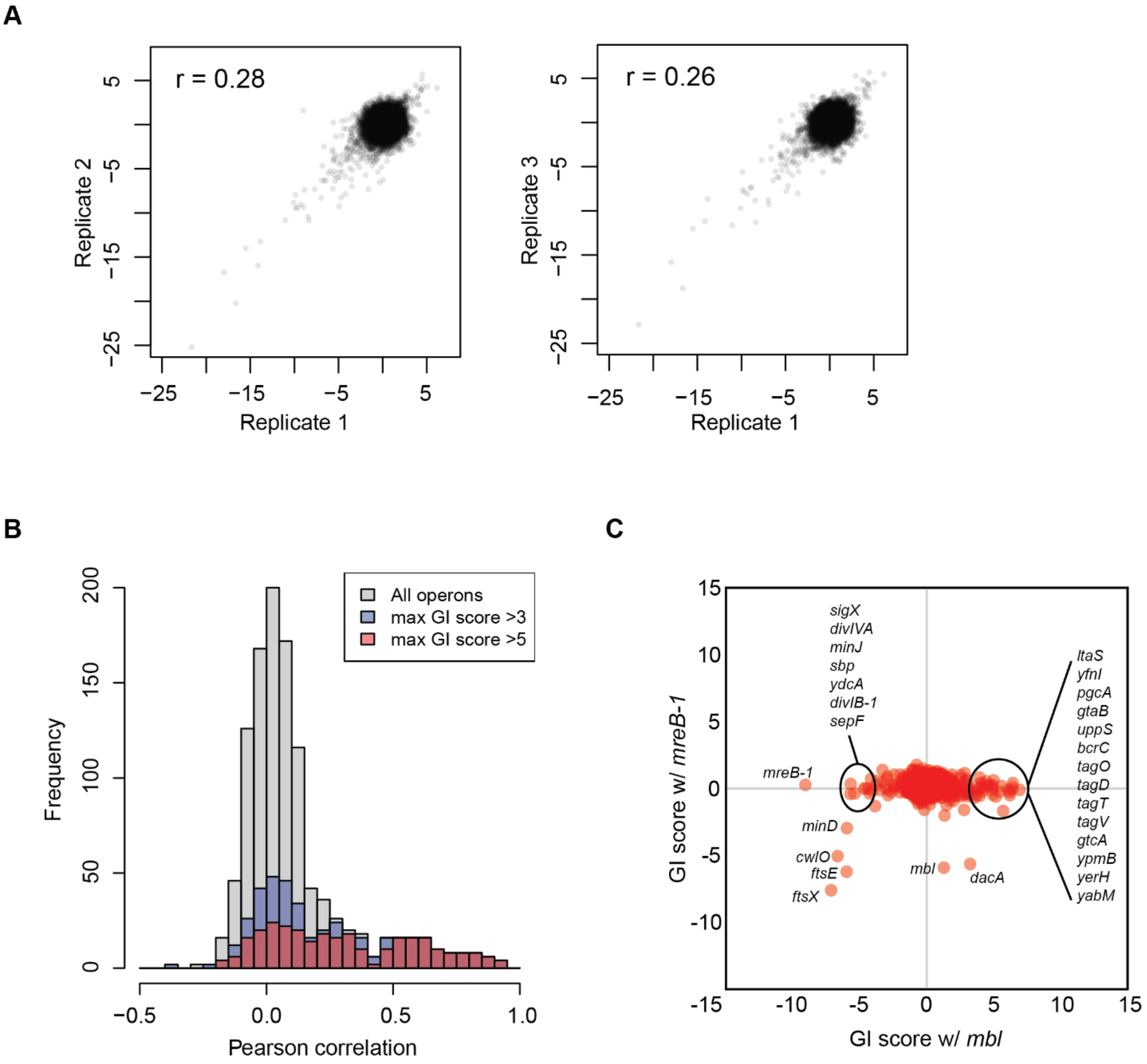
A
r = 0.28
Replicate 2
Replicate 1
r = 0.26
Replicate 3
Replicate 1
−25
−15
−5
5
B
All operons
max GI score >3
max GI score >5
Frequency
Pearson correlation
0
50
100
150
200
−0.5
0.0
0.5
1.0
C
GI score w/ *mreB-1*
GI score w/ *mbl*
-15
-10
-5
0
5
10
15
*sigX*
*divIVA*
*minJ*
*sbp*
*ydcA*
*divIB-1*
*sepF*
*mreB-1*
*minD*
*cwlO*
*ftsE*
*ftsX*
*mbl*
*dacA*
*ltaS*
*yfnI*
*pgcA*
*gtaB*
*uppS*
*bcrC*
*tagO*
*tagD*
*tagT*
*tagV*
*gtcA*
*ypmB*
*yerH*
*yabM*

**Figure S6**

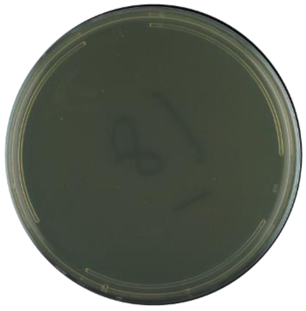

Δ*ltaS* Δ*dltABCDE*
16 hours

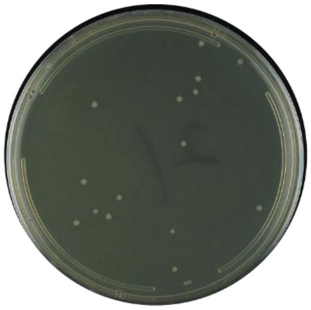

Δ*ltaS* Δ*amyE*
16 hours

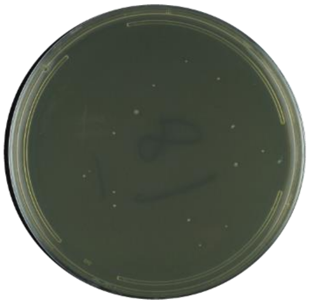

Δ*ltaS* Δ*dltABCDE*
40 hours

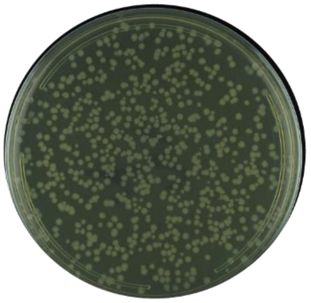

Δ*yfnI* Δ*dltABCDE*
16 hours

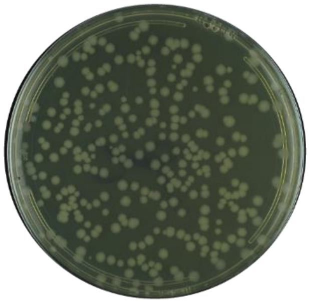

Δ*yfnI* Δ*amyE*
16 hours

**Figure S7**

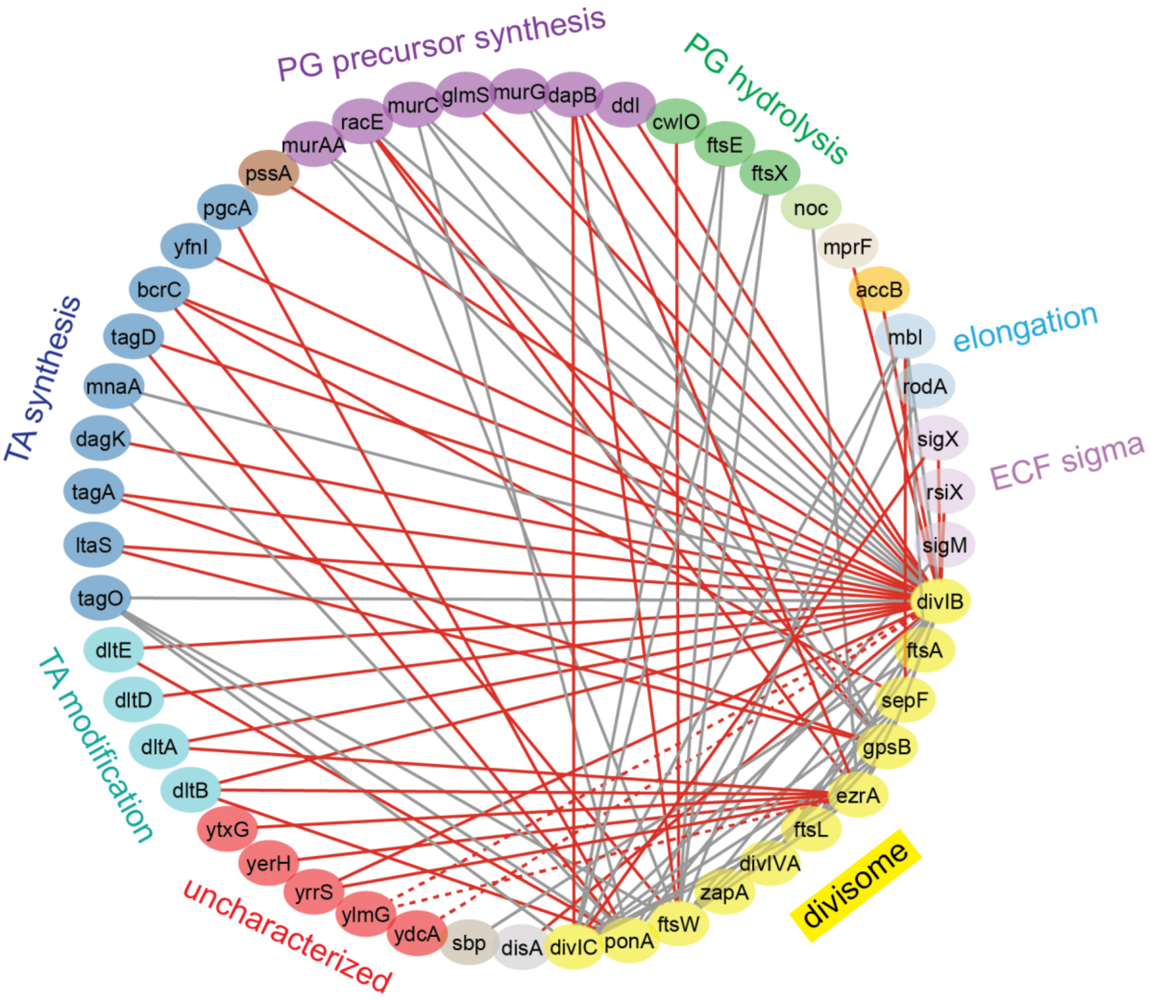
PG precursor synthesis
PG hydrolysis
elongation
ECF sigma
divisome
uncharacterized
TA modification
TA synthesis
murAA
racE
murC
glmS
murG
dapB
ddl
cwlO
ftsE
ftsX
noc
mprF
accB
mbl
rodA
sigX
rsiX
sigM
divIB
ftsA
sepF
gpsB
ezrA
ftsL
divIVA
zapA
ftsW
ponA
divIC
disA
sbp
ydcA
ylmG
yrrS
yerH
ytxG
dltB
dltA
dltD
dltE
tagO
ltaS
tagA
dagK
mnaA
tagD
bcrC
yfnI
pgcA
pssA

### Figure S8

**A**

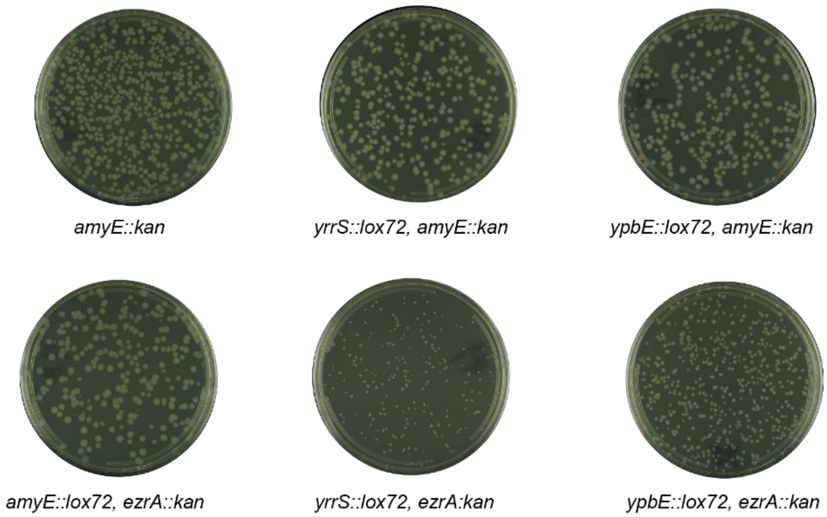
amyE::kan
yrrS::lox72, amyE::kan
ypbE::lox72, amyE::kan
amyE::lox72, ezrA::kan
yrrS::lox72, ezrA:kan
ypbE::lox72, ezrA::kan

**B**

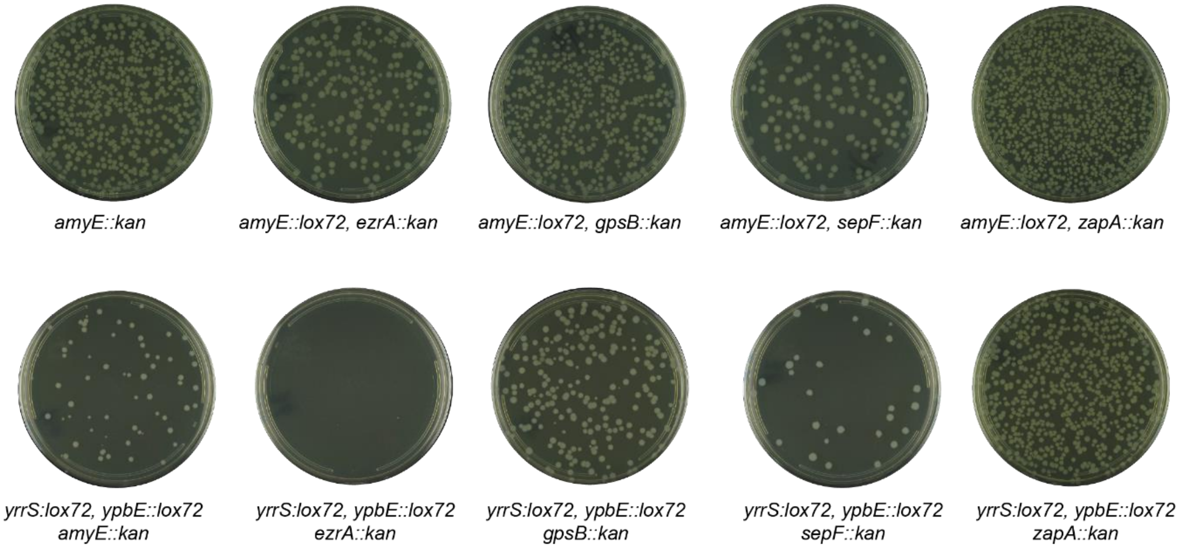
amyE::kan
amyE::lox72, ezrA::kan
amyE::lox72, gpsB::kan
amyE::lox72, sepF::kan
amyE::lox72, zapA::kan
yrrS:lox72, ypbE::lox72 amyE::kan
yrrS:lox72, ypbE::lox72 ezrA::kan
yrrS:lox72, ypbE::lox72 gpsB::kan
yrrS:lox72, ypbE::lox72 sepF::kan
yrrS:lox72, ypbE::lox72 zapA::kan
