## Supplementary Note for "Comprehensive double-mutant analysis of the *Bacillus subtilis* envelope using double-CRISPRi"

### Supplementary Note 1

#### Further characterization of the Δ*mbl*-suppressor mutants

Activation of the sigma factor SigI (and presumably upregulation of its operon) by deletion of the gene encoding its anti-sigma factor *rsgI* was previously shown to suppress the essentiality of *mbl* (Schirner and Errington, 2009). We therefore tested whether the *mbl* suppressors we isolated worked by activating SigI. Importantly, we were able to reconstruct many *mbl*-suppressor double-deletions in a Δ*sigI* background (Table N1). The triple mutant growth phenotypes were similar to those of the double mutants, suggesting that these suppressive mechanisms are SigI-independent.

We observed that many of our *mbl*-suppressor double-deletion strains, as well as the triple-deletion strains harboring Δ*sigI* lysed after overnight growth. This observation is consistent with the less positive GI scores of the *mbl*-suppressor strains following overnight growth observed in the double-CRISPRi screen (Table S3), and was also apparent on LB agar plates as lysed colonies after extended growth (Figures N1A and N1B). To identify mutations that further stabilize the *mbl*-suppressor strains, we isolated 11 colonies that outgrew from lysed colonies (Figure N1C) and sequenced their genomes. Suppressors isolated in both the double- and triple-deletion strains mapped to: *walK,* the kinase component of the essential cell wall homeostasis regulator WalRK; its negative regulator *walH*; and *yhdK*, the negative regulator of the cell wall sigma factor SigM (Table N1) (Szurmant et al., 2005; Zhao et al., 2019). Mutations and truncations in negative regulators are commonly loss-of-function mutants. Therefore the observed mutations likely increase WalK sensor kinase activity or activate SigM, which is required for the regulation of PG synthesis. Thus, the suppressors identified in our screen partially compensate for the lack of Mbl during exponential growth, but further activation of PG synthesis systems is required for survival during stationary phase growth.

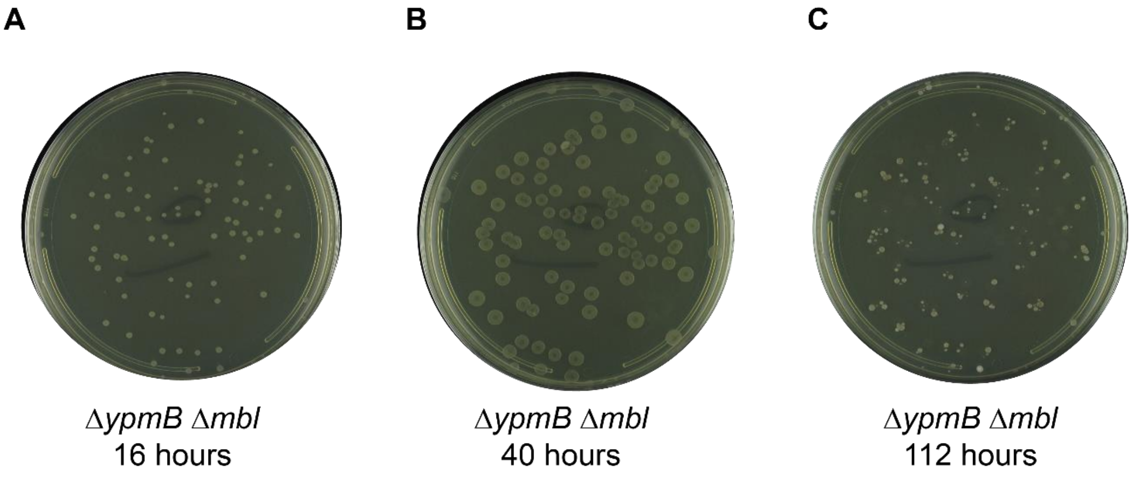


**Figure N1. The growth of mbl and suppressor double deletion mutant.** *mbl::kan* fragment was transformed into an antibiotic marker-free *ypmB* deletion strain and incubated for A) 16 hours, B) 40 hours, and C) 112 hours. We observed similar growth patterns of most of the other *mbl*/suppressor double deletion mutants on the LB agar plates.

**Table N1. List of the second mutations that might stabilize mbl/suppressor double deletion strains.**

| Strain | genotype | second mutations | | |
|---|---|---|---|---|
| | | mutation/position | gene | predicted phenotype |
| BKS0010 | *ltaS::lox72, mbl::kan* | L190V (TTG→GTG) | *walK* | WalK activation? |
| BKS0020 | *bcrC::lox72, mbl::kan* | Δ14 bp after 837/1368 nt | *walH* | WalK activation |
| BKS0030 | *ypmB::lox72, mbl::kan* | P391S (CCG→TCG) | *walK* | WalK activation? |
| BKS0040 | *ypmB::lox72, mbl::kan* | Δ2 bp after 655/1368 nt | *walH* | WalK activation |
| BKS0050 | *yvcK::lox72, mbl::kan* | L198F (CTC→TTC) | *walK* | WalK activation? |
| BKS0060 | *yvcK::lox72, mbl::kan* | S143N (AGT→AAT) | *sigM* | SigM activation? |
| BKS0070 | *sigI::lox72, yfnI::lox72, mbl::kan* | Δ21 bp after131/291 nt | *yhdK* | SigM activation |
| BKS0080 | *sigI::lox72, yfnI::lox72, mbl::kan* | V553G (GTG→GGG) | *walK* | WalK activation? |
| BKS0090 | *sigI::lox72, yvcK::lox72, mbl::erm* | Δ1 bp after 721/1368 nt | *walH* | WalK activation |
| BKS0100 | *sigI::lox72, yvcK::lox72, mbl::erm* | F244S (TTT→TCT) | *walK* | WalK activation? |
| BKS0110 | *sigI::lox72, yerH::lox72, mbl::kan* | duplication (CGGCAGCCATGGCG AACGGAA) at 131/291 nt | *yhdK* | SigM activation |

### Supplementary Note 2

### GI profile of *ydcA*, an uncharacterized gene

In our screen *ydcA* GIs were highly correlated with those of the *dlt* operon and *sigX* (Figure 3C and Table N2). *ydcA* has strong negative GIs with *pgcA*, *gtaB*, *divIB*, *mbl*, *prsW*, *cwlO*, and *ggaA*. As expected, these negative GIs (except for *mbl* and *prsW)* were shared with the *dlt* operon and *sigX* (Table N3). To better understand these interactions, we constructed two pairs of double deletion mutants (*pgcA/ydcA* and *gtaB/ydcA*) using pairwise donor-recipient combinations in transformations (*pgcA::lox72/ydcA::kan*, *ydcA::lox72/pgcA::kan*, *gtaB::lox72/ydcA::kan*, *ydcA::lox72/gtaB::kan*). Interestingly, only the mutants harboring the kanamycin-resistance gene replacing *ydcA* recapitulated the negative GIs, suggesting that the kanamycin-resistance gene at *ydcA* rather than the *ydcA* deletion itself was the major contributor to the observed phenotypes.

This finding prompted us to examine the genomic context of the *ydcA* gene locus. We found that the almost the entirety of the *ydcA* gene overlapped the 5'UTR of *acpS*, which encodes the essential acyl-carrier protein synthetase and is transcribed in the opposite direction from *ydcA* (Figure N2). Given this genomic arrangement, two explanations for the GIs of *ydcA* are possible. First and most likely, CRISPRi knockdown of *ydcA* or replacement of *ydcA* with the kanamycin-resistant gene may reduce the expression of *acpS* by targeting the template strand of its promoter with CRISPRi or by producing antisense RNA for 5'UTR of *acpS* via strong transcription of the kanamycin-resistance gene. AcpS interacts with DltC, and this interaction is essential for D-alanylation of TA (Ma et al., 2018; Nikolopoulos et al., 2022). This implies that *acpS* knockdown partially produces the phenotype of *dlt* genes knockdown. Consistent with the idea that the phenotypes of *ydcA* are due to downregulation of *acpS*, *ydcA* is a member of SigE regulon and therefore primarily expressed during sporulation rather than under normal growth conditions such as LB (Eichenberger et al., 2003). Together, these data suggest that the phenotypes of the *ydcA* knockdown strain result from the partial knockdown of *acpS* followed by reduced D-alanylation of TA. A second, though less likely, explanation for the phenotype is that reduced expression of *ydcA* itself produced GI with other genes. A *ydcA* homolog in *S. aureus*, *actH* was identified as an activator of LytH, an amidase involved in cell growth and division (Do et al., 2020). Thus, *ydcA* may contribute to cell wall homeostasis by modulating certain cell wall hydrolase activity even though we didn't observe any phenotype for *yqiI*, a homolog of S. aureus *lytH*.

Table N2. Highly correlated GI with *ydcA* (Pearson's r >0.5)

| gene | correlation with *ydcA* |
|---|---|
| *sigX* | 0.84 |
| *dltE* | 0.73 |
| *dltA* | 0.69 |
| *dltB* | 0.68 |
| *dltD* | 0.67 |
| *rsiX* | 0.57 |

Table N3. GI profile of *ydcA* (|GI|>2.5) and highly correlated genes

| sgRNA | GI_*ydcA* | GI_*sigX* | GI_*dltE* | GI_*dltA* | GI_*dltB* | GI_*dltD* | GI_*rsiX* |
|---|---|---|---|---|---|---|---|
| *pgcA* | -10.23 | -10.84 | | | -5.70 | -5.44 | |
| *gtaB* | -10.11 | -10.43 | -10.74 | -7.86 | -6.89 | -3.90 | |
| *divIB-1* | -9.07 | -8.80 | -3.01 | -3.60 | -3.66 | -3.37 | -5.61 |
| *divIB-2* | -6.61 | -9.92 | -4.98 | -5.81 | -4.09 | -5.27 | |
| *mbl* | -4.54 | -5.71 | -2.65 | -0.80 | -0.84 | -0.74 | |
| *prsW* | -2.84 | 0.24 | -1.19 | 0.90 | 0.11 | 0.45 | |
| *cwlO* | -2.80 | -2.65 | -4.51 | -3.17 | -1.96 | -3.48 | -2.51 |
| *ggaA* | -2.72 | -3.01 | -4.46 | -3.06 | -2.92 | -3.85 | -1.13 |

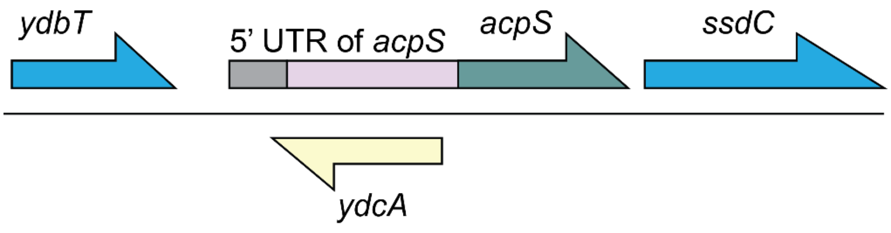


**Figure N2. Genomic context of *ydcA* locus**

### Supplementary Note 3

#### Additional GI data support a role for *yrrS*, *ypbE, ytxG*, and *yerH* in cell division

Our newly identified cell division genes all exhibited negative GIs with *ezrA*, but also had distinct GI, as evidenced by the lack of significant GI correlations among them (Table S3 and S4).

Like *ezrA* and *gpsB,* both *yrrS* and *ytxG* exhibited negative GIs with *ponA* (encodes PBP1), raising the possibility that they, like EzrA and GpsB, play a role in PBP1 localization. As YpbE physically interacts with YrrS and has a similar protein-protein interaction profile to that of YrrS (Cleverley et al., 2019), it may also be involved in this process. These proteins may have additional roles in division as there are no significant GI correlations among them (Table S4). For example, *yrrS* has a strong negative GI with *divIB* whereas *ytxG* has many negative GIs with other genes including *mreB, uppS, and bcrC*. Interestingly, as mentioned above, *yerH* suppressed the *mbl* phenotype, suggesting that this gene might be involved in a connection between cell elongation and division independent of GpsB. In summary, our GI data and morphological analysis data support the idea that we have identified new players in cell division.

### Supplementary Note 4

### Disruption of newly identified cell division genes results in membrane and cell-shape defects

In addition to filamentation and other gross phenotypes, Δ*yrrS*, Δ*yerH*, Δ*ypbE,* and Δ*gpsB* all exhibited an increased number of FM4-64 foci, suggesting membrane accumulation (Figure N3A). These foci occurred 20-40% of cells in the Δ*yrrS*, Δ*yerH*, Δ*ypbE,* and Δ*gpsB* strains, and only ~10% of unperturbed cells (*ezrA-KD,* knockdown not induced; Figure N3B). Membrane foci have been previously associated with membrane invaginations due to incorrect recruitment of septal factors (Bartlett et al., 2024; Gao et al., 2017), suggesting that the Δ*yrrS*, Δ*yerH*, Δ*ypbE,* and Δ*gpsB* strains have pre-existing defects in PBP localization that are exacerbated upon *ezrA* knockdown. This idea is supported by time lapse imaging showing that these foci frequently nucleate morphological defects such as bending or lysis (Fig. N3A).

In some cells, apparently fully formed membrane cross-bands and invaginations disappeared (Figure N3C). In such events, the cell body sometimes swelled uniformly nearby (Figure N3C, knockdown of *ezrA* in Δ*gpsB*) or developed a sharp kink (Fig. N3B, knockdown of *ezrA* in Δ*yrrS*) indicative of asymmetric growth around the circumference. This behavior occurred in multiple strains, and is consistent with previous findings that an *ezrA gpsB* double mutant exhibits bulging due to a failure in completing cell pole maturation (Claessen et al., 2008). As previously reported (Bartlett et al., 2024), Δ*ytxG* cells exhibited bright membrane patches at the cell poles (Figure N3A, blue arrows) as well as along the cell body (Figure N3A, Δ*ytxG*, red arrows), the latter potentially due to membrane invagination. Collectively, these data suggest that specific gene knockouts combined with *ezrA* depletion can form functional Z-rings but are defective in synthesizing cell poles at Z-ring locations.

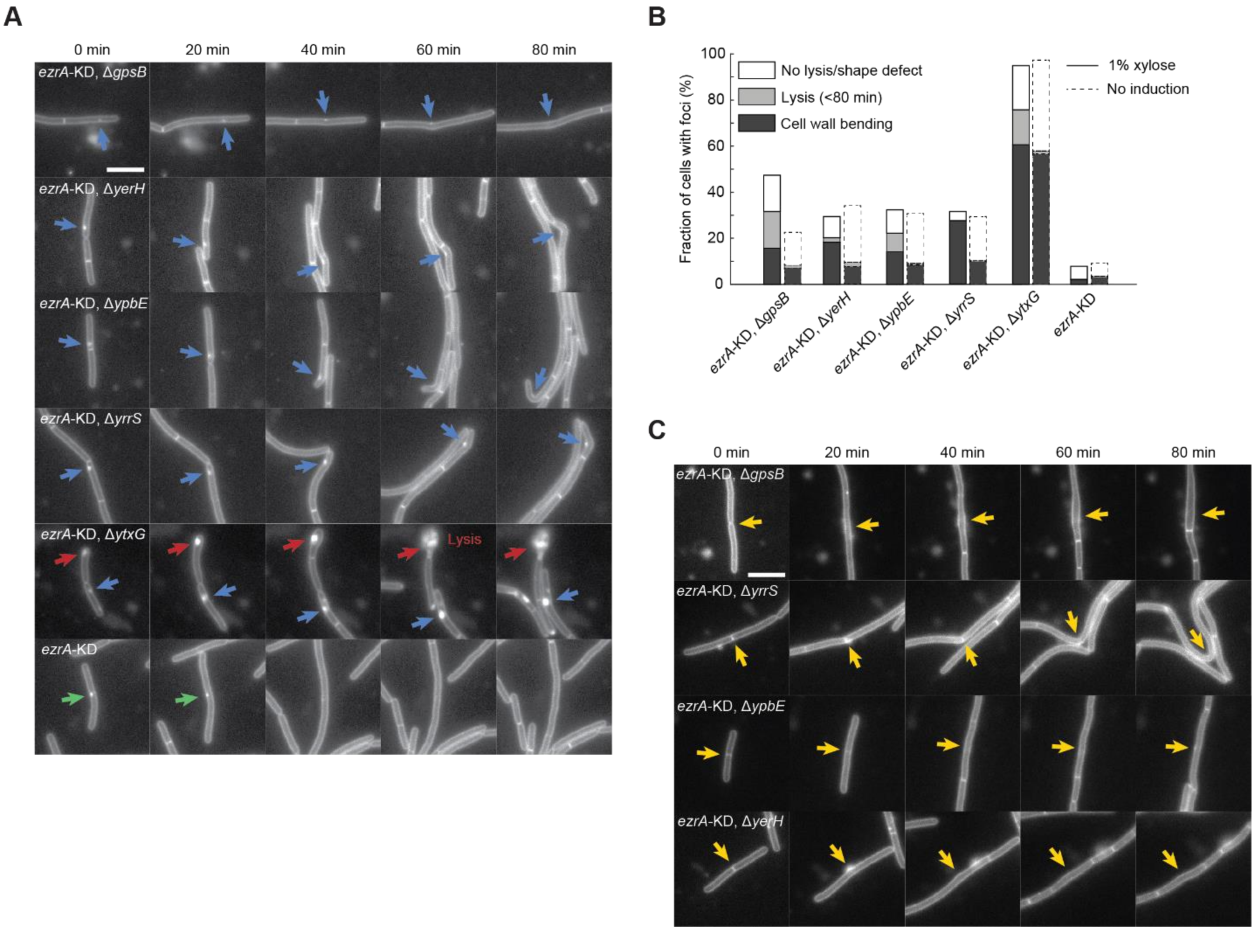


**Figure N3. Knockout mutants of several uncharacterized genes exhibit distinct shapes and membrane phenotypes under *ezrA* depletion. A)** Cells with bright foci along the cell body can develop sharp kinks due to asymmetric cell wall growth (blue arrows). Δ*ytxG* cells with bright patches around cell poles can exhibit bulging poles and/or lysis (red arrows). Without additional gene knockouts, bright foci under *ezrA* depletion typically does not cause shape defects. Scale bar: 5 µm. **B)** The fraction of cells containing bright foci and those developing shape defects or undergoing lysis is dependent on the deletion of specific genes and the depletion of EzrA, underscoring the impact of genetic interactions on cell division defects. **C)** Examples of cell poles or FM4-64 cross-bands that disappear after formation (red arrows). In some cases, swelling (Δ*gpsB*) or bending (Δ*yrrS* and Δ*yerH*) occurs at the location of the disappearing pole/cross-band during subsequent growth of the cell wall. Arrows indicate the poles/cross-bands or the subsequent shape defect. Scale bar: 5 µm.

### Supplemental Note References


Bartlett, T.M., Sisley, T.A., Mychack, A., Walker, S., Baker, R.W., Rudner, D.Z., and Bernhardt, T.G. (2024). FacZ is a GpsB-interacting protein that prevents aberrant division-site placement in Staphylococcus aureus. Nat Microbiol *9*, 801-813.

Claessen, D., Emmins, R., Hamoen, L.W., Daniel, R.A., Errington, J., and Edwards, D.H. (2008). Control of the cell elongation-division cycle by shuttling of PBP1 protein in Bacillus subtilis. Mol Microbiol *68*, 1029-1046.

Cleverley, R.M., Rutter, Z.J., Rismondo, J., Corona, F., Tsui, H.T., Alatawi, F.A., Daniel, R.A., Halbedel, S., Massidda, O., Winkler, M.E., *et al.* (2019). The cell cycle regulator GpsB functions as cytosolic adaptor for multiple cell wall enzymes. Nature communications *10*, 261.

Do, T., Schaefer, K., Santiago, A.G., Coe, K.A., Fernandes, P.B., Kahne, D., Pinho, M.G., and Walker, S. (2020). Staphylococcus aureus cell growth and division are regulated by an amidase that trims peptides from uncrosslinked peptidoglycan. Nat Microbiol *5*, 291-303.

Eichenberger, P., Jensen, S.T., Conlon, E.M., van Ooij, C., Silvaggi, J., Gonzalez-Pastor, J.E., Fujita, M., Ben-Yehuda, S., Stragier, P., Liu, J.S., *et al.* (2003). The sigmaE regulon and the identification of additional sporulation genes in Bacillus subtilis. J Mol Biol *327*, 945-972.

Gao, Y., Wenzel, M., Jonker, M.J., and Hamoen, L.W. (2017). Free SepF interferes with recruitment of late cell division proteins. Sci Rep *7*, 16928.

Ma, D., Wang, Z., Merrikh, C.N., Lang, K.S., Lu, P., Li, X., Merrikh, H., Rao, Z., and Xu, W. (2018). Crystal structure of a membrane-bound O-acyltransferase. Nature *562*, 286-290.

Nikolopoulos, N., Matos, R.C., Courtin, P., Ayala, I., Akherraz, H., Simorre, J.P., Chapot-Chartier, M.P., Leulier, F., Ravaud, S., and Grangeasse, C. (2022). DltC acts as an interaction hub for AcpS, DltA and DltB in the teichoic acid D-alanylation pathway of Lactiplantibacillus plantarum. Sci Rep *12*, 13133.

Schirner, K., and Errington, J. (2009). The cell wall regulator sigmaI specifically suppresses the lethal phenotype of mbl mutants in Bacillus subtilis. J Bacteriol *191*, 1404-1413.

Szurmant, H., Nelson, K., Kim, E.J., Perego, M., and Hoch, J.A. (2005). YycH regulates the activity of the essential YycFG two-component system in Bacillus subtilis. J Bacteriol *187*, 5419-5426.

Zhao, H., Roistacher, D.M., and Helmann, J.D. (2019). Deciphering the essentiality and function of the anti-sigma(M) factors in Bacillus subtilis. Mol Microbiol *112*, 482-497.
